# A light-induced microprotein triggers regulated intramembrane proteolysis to promote photo sensing in a pathogenic bacterium

**DOI:** 10.1101/2025.04.28.651061

**Authors:** Dimitrios Manias, Jonathan Winkelman, Gong Chen, Sampriti Mukherjee

## Abstract

Light, a ubiquitous environmental stimulus, shapes behavior and physiology across all domains of life. While photoreceptors are widespread in bacterial genomes, their functional roles and signal transduction mechanisms in non-photosynthetic bacteria remain understudied. Light represses biofilms and virulence factors through a bacteriophytochrome photoreceptor BphP and response regulator AlgB in the human pathogen *Pseudomonas aeruginosa*. Here, we used transposon mutagenesis screening to identify a conserved hypothetical microprotein, DimA, as the master activator of the photo-sensing cascade. Transcriptomics, luciferase reporter assays and physiological assays revealed that deletion of *dimA* abolishes light-dependent suppression of virulence factors and biofilms. Mechanistically, we demonstrated that DimA activates the site-I protease AlgW, triggering regulated intramembrane proteolysis of the anti-sigma factor MucA, liberating sigma factor AlgU (σ22), which promotes *algB* expression. We discovered a positive feedback loop where light-activated AlgB upregulates *dimA* expression, thereby amplifying the photosensory response. This work establishes DimA as a crucial activator of photo sensing and expands our understanding of bacterial adaptation to changing light environments.

## INTRODUCTION

Bacteria employ sophisticated molecular mechanisms to perceive and respond to environmental stimuli through specialized sensory apparatuses that are crucial for survival, pathogenesis, and ecological interactions, allowing bacteria to thrive in diverse and dynamic environments. Amongst primary sensing mechanisms are two-component signal transduction systems (TCS), comprising histidine kinases and response regulators, which detect various physicochemical parameters and initiate adaptive responses ([1], [2], [3]). Environmental perception encompasses chemoreception via membrane-bound receptors that recognize specific molecules, mechanosensation, cell-to-cell chemical communication via quorum sensing and photoreception via photoreceptor proteins such as bacteriophytochromes ([4], [5], [6], [7], [8]). Signal transduction cascades convert these environmental inputs into cellular responses through complex molecular networks. These include phosphorylation cascades, second messenger systems (particularly cyclic nucleotides like cAMP and c-di-GMP), and modulation of sigma factors and transcriptional regulators ([9], [10], [11], [12], [13]). These pathways culminate in the activation of specific genetic programs, enabling bacteria to alter metabolism, virulence factor production, biofilm formation, and motility accordingly.

Light, a ubiquitous environmental stimulus, is detected by photoreceptors in all domains of life ([14]). Particular photoreceptor photosensory domains are activated by specific wavelengths of light ([15]), with bacteriophytochromes being the most abundant photoreceptors ([16], [17]). Although photoreceptors are abundant in bacterial genomes, the molecular underpinnings of photo-sensing signal transduction in non-photosynthetic bacteria remain poorly understood, often due to the lack of a known phenotypic output. We previously reported that the human pathogen *Pseudomonas aeruginosa* senses and responds to light to inhibit biofilm formation and virulence factor production ([18]). The photo-sensing signaling cascade in *P. aeruginosa* is initiated by the activation of the biliverdin-containing bacteriophytochrome photoreceptor BphP by light. BphP is a histidine kinase that, in response to light, phosphorylates and activates the response regulator AlgB. Phospho-AlgB in turn represses the expression of biofilm and virulence genes. While this BphP-AlgB photo sensing two-component system is conserved in diverse bacterial phyla, relatively little is known about the factors that control photo sensing. One negative regulator of photo sensing is the phosphatase KinB that dephosphorylates AlgB to dampen photo-sensing response ([18], [19]). Previous studies on transcriptional regulation of *bphP* have implicated the stationary phase sigma factor RpoS ([20]). The expression of *algB* is promoted by the alternative sigma factor σ22 (also known as AlgT/AlgU) and in turn by regulated intramembrane proteolysis (RIP) ([21], [22], [23], [24], [25], [26], [27]). RIP is a widely conserved signal transduction mechanism involving sequential protein cleavage by Site-1 (S1P) and Site-2 (S2P) proteases ([28], [29], [30], [31], [32]). In *P. aeruginosa*, the anti-sigma factor MucA, the homolog of *Escherichia coli* RseA, is a signaling protein with a periplasmic domain, a single transmembrane helix and an N-terminal domain that binds and antagonizes AlgU ([33], [34]). MucA is sequentially cleaved via RIP by AlgW (S1P) and MucP (S2P). Following RIP, MucA is degraded by cytoplasmic ClpXP proteases and AlgU is released, leading to activation of *algB* expression.

In this work, we performed genome-wide transposon mutagenesis screening to identify positive regulators of photo sensing in *P. aeruginosa*. We discovered that a previously uncharacterized microprotein, PA14_20480, that we name DimA (*<u>D</u>iaphotos Initiator <u>M</u>icroprotein of <u>A</u>lg<u>B</u>*) positively regulated photo-sensing driven repression of biofilm formation and virulence factor production. DimA, a periplasmic protein, activated the site-I protease AlgW, triggering proteolysis of the anti-sigma factor MucA, and releasing the alternate sigma factor AlgU in turn to promote *algB* expression. We further identified a positive feedback loop whereby AlgB transcription factor drives *dimA* expression under far-red light, reinforcing the photo-sensing response.

## RESULTS

### DimA promotes photo sensing

We previously discovered that *P. aeruginosa* responds to light via the BphP-AlgB two-component system that represses biofilm formation and virulence factor production in the presence of light (Fig. 1a,b) ([18]). Specifically, far-red light inhibits the expression of the major biofilm matrix exopolysaccharide Pel in wildtype (WT) *P*. *aeruginosa* UCBPP-PA14 (hereafter called PA14) and virulence factors such as pyocyanin, hydrogen cyanide and lectin A ([18]). To discover regulators of the BphP-AlgB photo-sensing pathway, we first generated a light responsive reporter system by engineering a chromosomal fusion of the promoter of the gene encoding lectin A to a promoter-less luciferase cassette (*attB::PlecA’-‘luxCDABE,* hereafter referred to as *lecA-lux*) (Fig. S1). As expected, upon shining far-red light this reporter strain exhibits approximately ten-fold lower luminescence compared to dark condition (Fig 1c). Next, we screened ∼35,000 transposon insertion mutants for colonies exhibiting a “bright” phenotype, i.e., increased luminescence over the parent strain (Fig. S1) and identified 17 genes that had multiple independent insertions (Table S1). These can be broadly classified into three categories: (1) known photo-sensing components (*bphP* and *algB*), (2) known factors that control the transcription of *algB* (*algU*, *clpX*, *clpP*, *algW*, and *mucP*), and (3) hypothetical proteins. Category 1 hits serve as direct photoreceptor-response regulator (BphP-AlgB) TCS that can detect and respond to specific wavelengths of light, while Category 2 factors (AlgU, ClpX, ClpP, AlgW, and MucP) form the RIP signaling cascade that controls *algB* expression (Fig. 1d, S1, S2). Accordingly, Western blot analysis using custom raised antibodies against AlgB showed that AlgB protein levels were significantly reduced in the Category 2 transposon mutants compared to the single mutant of *ΔkinB* as well as WT PA14 (Fig. S2).

**Fig. 1:**
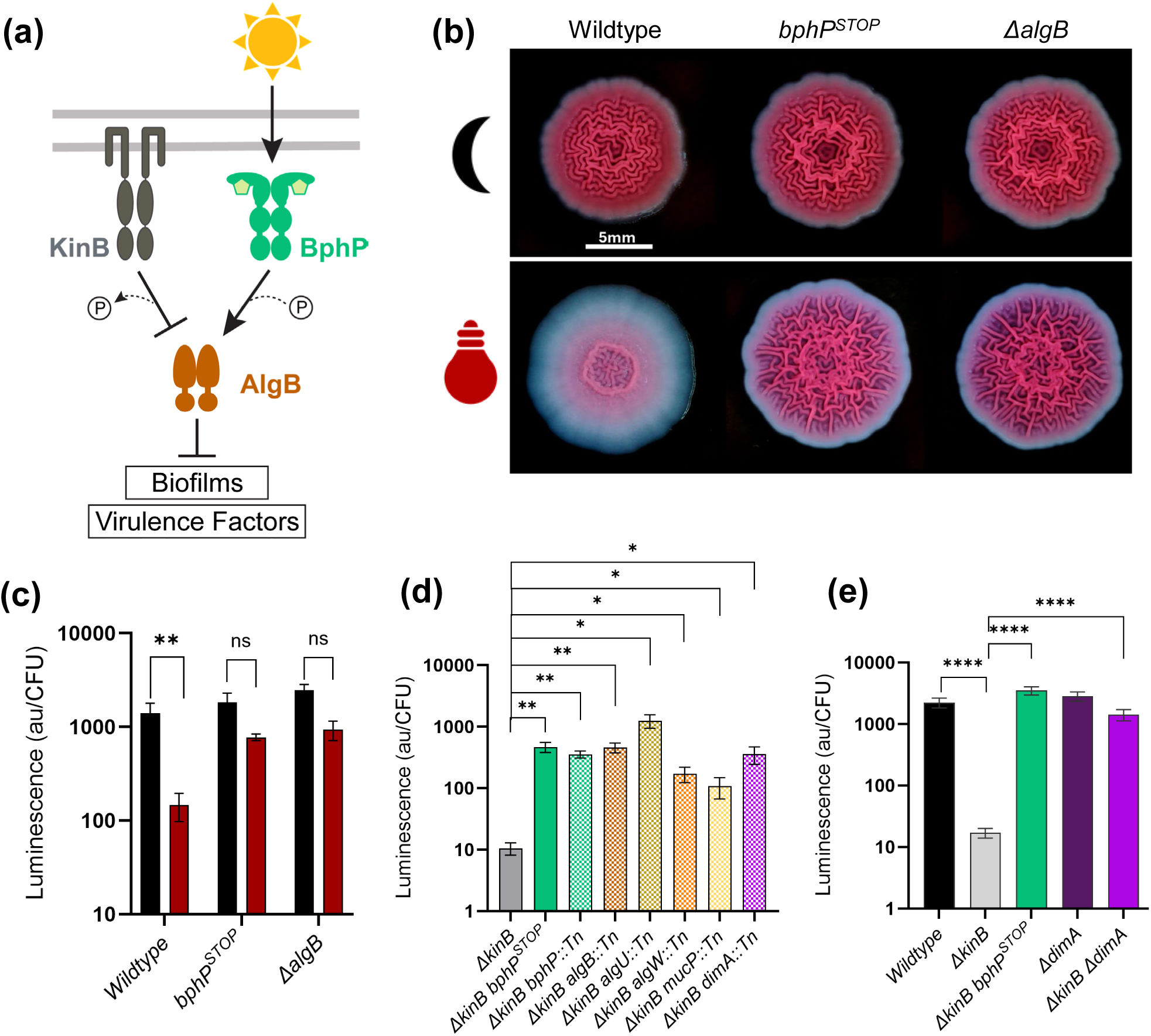
A genetic screen identifies positive regulators of photo sensing. (a) Schematic of BphP-AlgB photo-sensing signaling cascade in *P. aeruginosa*. Light (denoted by “sun” symbol) stimulates the BphP kinase (green) to auto-phosphorylate and subsequently transfer the phosphoryl group to AlgB (brown) to activate AlgB. AlgB∼P activates transcription of genes required for repression of group behaviors such as biofilm formation and virulence factor production. KinB (gray) acts as a phosphatase to antagonize AlgB. A “P” in a circle denotes addition or removal of a phosphate moiety. Arrow indicates activation and T-bar indicates inhibition. (b) Colony biofilms under dark (denoted by “black crescent” symbol) and far-red (denoted by “red bulb” symbol) for WT, *bphP^STOP^*, *ΔalgB*. Scale bar, 5 mm. (c) Quantification of *lecA-lux* activity in WT and mutant backgrounds under dark (black bars) and far-red light (red bars) conditions. Error bars represent SEM of three biological replicates. Statistical significance was determined using t-test pairwise comparisons in GraphPad Prism software. ** P <0.005 (d) Quantification of *lecA-lux* activity for Tn5-candidates. Error bars represent SEM of three biological replicates. Only pairwise comparisons that had P value <0.05 are denoted. Statistical significance was determined using t-test pairwise comparisons in GraphPad Prism software. ** P <0.005, * P <0.05 (e) Quantification of *lecA-lux* activity in WT and mutant backgrounds under ambient light conditions. Error bars represent SEM of three biological replicates. Only pairwise comparisons that had P value <0.05 are denoted. Statistical significance was determined using t-test pairwise comparisons in GraphPad Prism software. **** P <0.0001

Here, we focus on one transposon hit in a hypothetical gene, *PA14_20840*, that exhibited increased luminescence similar to those exhibited by the *bphP::Tn* and *algB::Tn* insertion mutants (Fig. 1d, S1). The gene *PA14_20480* encodes for a 92 amino acid microprotein (defined as proteins between 50 and 100 amino acids, [35]) that we refer to as DimA. To verify that DimA plays a role in *lecA* expression, we generated an in-frame marker-less deletion of *dimA* in the WT and *ΔkinB* strains bearing the *lecA-lux* reporter. Under ambient light conditions, the WT PA14, *bphP^STOP^*and *ΔkinB bphP^STOP^* strains exhibited strong luminescence signal while the absence of the negative regulator of photo sensing KinB reduced *lecA-lux* reporter expression about 130-folds compared to WT (Fig. 1e). Consistent with the transposon mutant result, the *ΔkinB ΔdimA* double mutant was unable to abolish *lecA* expression similar to the *ΔkinB bphP^STOP^* double mutant (Fig. 1e) suggesting that DimA is an inhibitor of Lectin A production.

DimA could function as a specific inhibitor of *lecA* expression or be a general photo-sensing regulator. To distinguish between these, we reasoned that a positive regulator of photo sensing must repress other behaviors inhibited by light such as pyocyanin production and biofilm formation. Accordingly, we measured the production of the phenazine pyocyanin. Similar to the *lecA-lux* reporter, pyocyanin production was not repressed in WT or the *ΔdimA* single mutant under ambient light conditions (Fig. 2a). However, the *ΔkinB* mutant had 75% lower pyocyanin levels compared to WT while the absence of DimA restored pyocyanin production (Fig. 2a). Complementation of the *ΔkinB ΔdimA* double mutant by introducing *dimA* under its native promoter in the *attB* chromosomal location reduced pyocyanin production by 72% (Fig. 2a), thereby demonstrating that DimA is a negative regulator of pyocyanin biosynthesis.

**Fig. 2:**
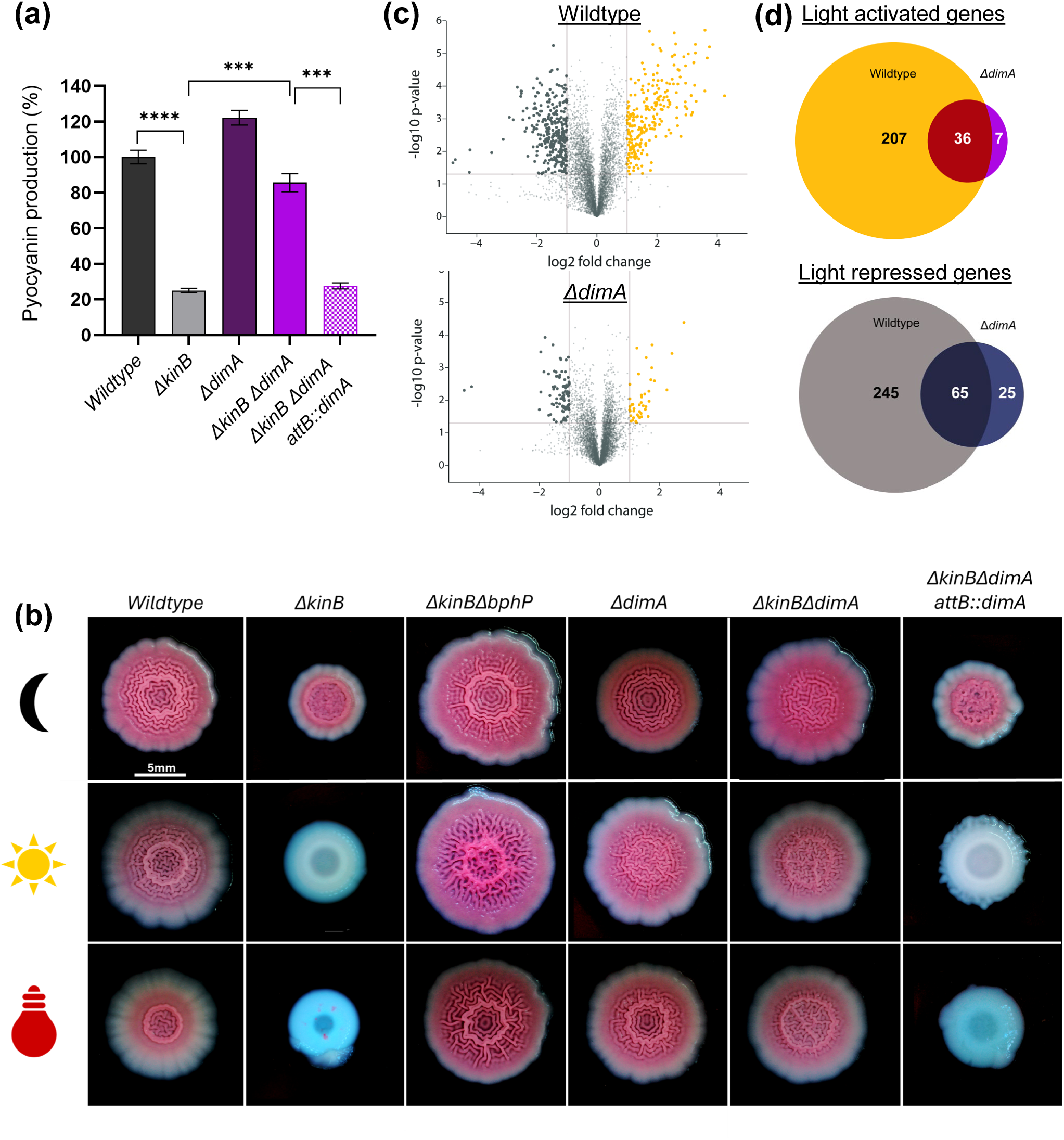
DimA promotes photo-sensing dependent behaviors. (a) Pyocyanin production (OD_695_) was measured and normalized to growth (OD_600_) in WT PA14 and the designated mutants grown overnight under ambient light conditions. Pyocyanin levels in WT PA14 were set to 100%. Error bars represent SEM of three biological replicates. Only pairwise comparisons that had P value <0.05 are denoted. Statistical significance was determined using t-test pairwise comparisons in GraphPad Prism software. *** P <0.001, **** P <0.0001. (b) Colony biofilms under dark (denoted by “black crescent” symbol), ambient (denoted by “sun” symbol) and far-red (denoted by “red bulb” symbol) for WT and mutant strains. Scale bar, 5 mm. (c) Volcano plots of RNA-seq data for WT and *ΔdimA* in the presence or absence of far-red light. Dark gray solid circles represent genes that are significantly repressed by far-red light by >2-fold while mustard yellow solid circles represent genes that are significantly activated by far-red light >2-fold. Light gray solid circles represent genes with gene expression fold changes ≥-2 or ≤2 or Padj ≥ 0.05. (d) Venn diagrams showing overlaps in genes that are differentially regulated in the presence and absence of light between WT and *ΔdimA* mutant. Numbers indicate number genes differentially regulated in each intersection.

Along with repression of virulence factors, the AlgB-BphP photo-sensing system regulates biofilm formation (Fig. 1b) ([18]). To determine whether DimA regulates light-mediated repression of biofilms, we assayed our strains for colony biofilm formation on Congo Red plates under light and dark conditions. In the absence of light, all the strains formed a rugose colony pattern indicative of biofilm formation (Fig. 2b). Exposure to ambient light, resulted in smooth colony pattern and repression of biofilm formation for the single mutant of *ΔkinB* (Fig. 2b). Absence of the photoreceptor BphP or DimA in the *ΔkinB* background showed de-repression of biofilms formation i.e., rugose colony phenotypes (Fig. 2b). The complemented strain *ΔkinB ΔdimA attB::P_dimA_-dimA* restored its ability to repress biofilms similar to the *ΔkinB* strain (Fig. 2b). We conclude that DimA is a repressor of biofilm formation.

To define the extent of DimA’s role in photo sensing, we used RNA sequencing to compare the genomic transcriptional profiles of the WT and *ΔdimA* strains. We performed the experiment under dark and far-red light conditions. Principal component analysis (PCA) of normalized read counts showed that samples of dark conditions clustered separately from those of far-red light (Fig. S3). Comparative transcriptomic analysis revealed a total of 553 differentially expressed genes (DEGs; genes with expression fold-changes ≤−2 and ≥2, and the P values (Padj), adjusted using the Benjamini-Hochberg procedure, <0.05) in the WT under far-red light compared to dark (Fig. 2c,d). These results confirm that far-red light drives large-scale transcriptional reprogramming in *P. aeruginosa*, controlling ∼10% of the genome. In the *ΔdimA* mutant, we find a total of 133 DEGs, of which 101 transcripts overlapped with the light regulon in WT. Thus, the *ΔdimA* mutant showed a 76% reduction in the number of DEGs compared to WT (Fig. 2c,d). Taken together, we conclude that DimA is a master activator of the photo-sensing response in *P. aeruginosa*.

### DimA positively regulates AlgB protein levels

One way in which DimA might promote photo sensing is by activating the expression of the photoreceptor BphP and/or the response regulator AlgB. Therefore, we determined the protein levels of BphP and AlgB in the WT and *ΔdimA* mutant strains. Western blot analysis using anti-FLAG antibody to probe for chromosomally encoded BphP-3xFLAG revealed that the protein levels of the photoreceptor BphP were not altered by the presence or absence of DimA (Fig. 3a,b). In contrast, Western blot analysis using custom raised antibodies against AlgB showed that the levels of the response regulator AlgB were significantly reduced in the *ΔkinB ΔdimA* double mutant compared to the single mutant of *ΔkinB* as well as WT PA14 (Fig. 3a,c). Furthermore, ectopic complementation of *dimA* in the *ΔkinB ΔdimA* double mutant restored AlgB levels comparable to the *ΔkinB* mutant indicating that DimA functions as an activator of AlgB protein (Fig. 3a,c). We conclude that while DimA does not alter BphP production, the absence of DimA significantly reduced AlgB protein levels.

**Fig. 3:**
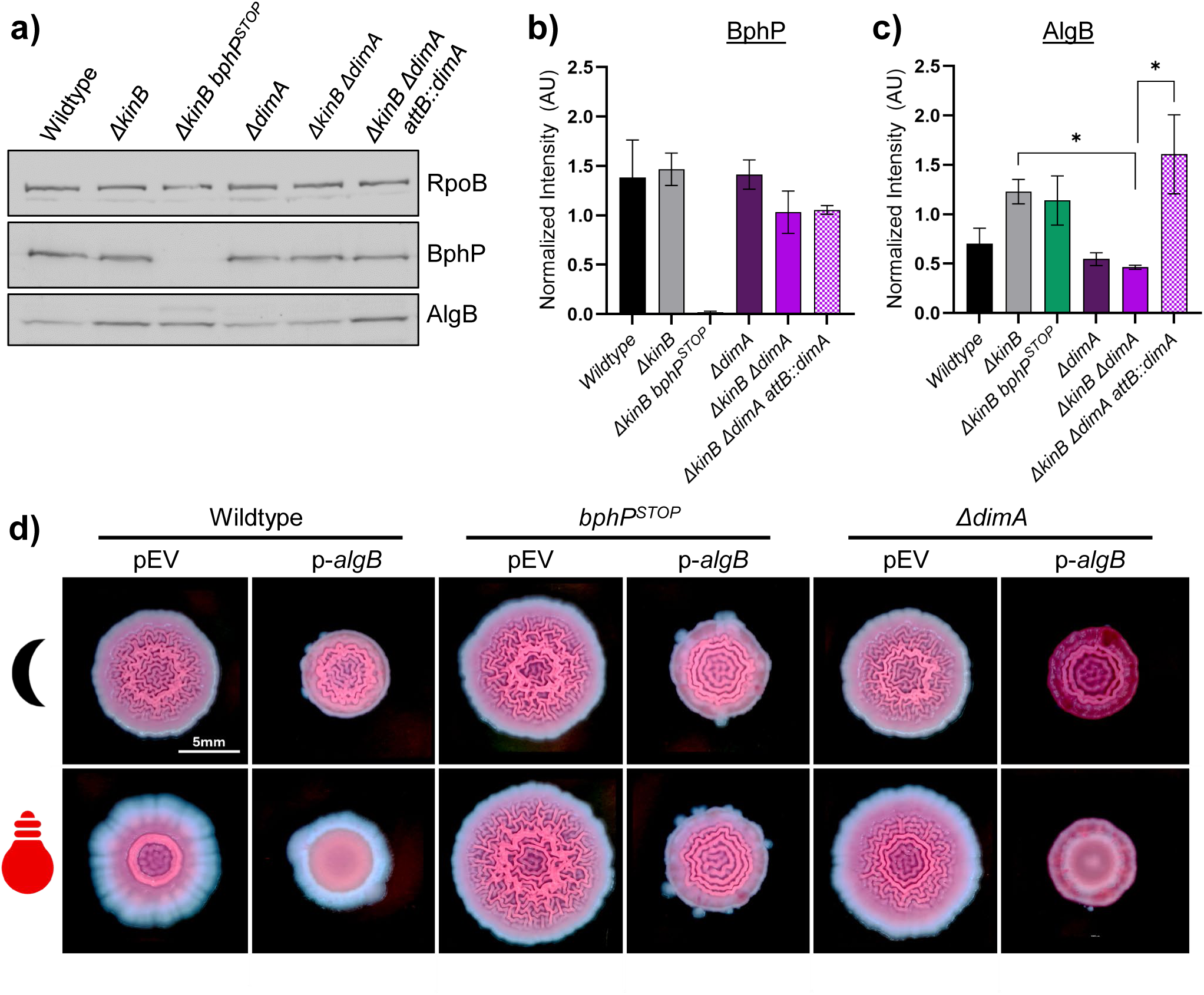
DimA enhances AlgB protein levels. (a) Western blot analysis of BphP-3xFLAG and AlgB protein levels. Overnight cultures of *P. aeruginosa* wildtype and the designated mutant strains were grown in triplicates in presence of ambient light. Representative blot of one of the replicates is shown, RpoB was used as a loading control. (b-c) Quantification of Western blots probing for BphP and AlgB. The values have been normalized to RpoB protein levels. Error bars represent SEM of three biological replicates. Statistical significance was determined using t-test pairwise comparisons in GraphPad Prism software. ** P <0.005, **** P <0.0001. (d) Colony biofilm phenotypes of WT PA14 and the designated mutants on Congo red agar medium supplemented with carbenicillin and grown for 72 h. Strains were grown under dark (Top row, denoted by “black crescent” symbol) and far-red (Bottom row, denoted by “red bulb” symbol). Strains are harboring empty vector of pUCP18 plasmid (pEV) or are harboring vector overexpressing AlgB protein (p*algB*). Scale bar is 5 mm for all images.

To define the genetic relationship between *algB* and *dimA,* we overexpressed *algB* on a multicopy plasmid and tested if that could compensate for the loss of DimA. Wildtype PA14 transformed with a vector containing *algB* under a constitutively active *P_lac_* promoter exhibited a reduction in biofilms when exposed to ambient light (Fig. 3d). Conversely, the biofilms were not repressed in the absence of the photoreceptor BphP even upon *algB* overexpression, as AlgB cannot get phosphorylated and therefore activated (Fig. 3d). However, overexpression of *algB* was sufficient to repress biofilms under light in the *ΔdimA* background, showing that AlgB’s function is epistatic to DimA (Fig. 3d). Taken together, we conclude that DimA is an upstream positive regulator of AlgB protein levels.

### The WTF C-terminal motif of DimA is essential for its function

To gain insight into DimA’s mode of action, we questioned whether DimA is present in other bacteria. Examination of over ninety complete *P. aeruginosa* genomes identified DimA orthologues in both clinical and environmental isolates (Fig. 4a, S4). In addition, reciprocal BLAST identified DimA homologs in other Pseudomonads such as *Pseudomonas syringae* and *Pseudomonas chlororaphis* (Fig. 4a, S4). The amino acid sequences of DimA orthologues are highly conserved (Fig. S5), suggesting that DimA has a conserved function. Next, we examined the predicted AlphaFold structure of the protein (Fig. 4b). Although the structure showed a very low average of per-residue model confidence score (pLDDT=49.03), it was predicted to have an N-terminal α-helix that could function as a signal peptide for Sec secretion system (Fig. 4c). To define the localization of DimA we engineered a C-terminal translational fusion of DimA with mNeonGreen fluorescent protein under IPTG inducible promoter using the plasmid pME6032. Upon IPTG induction, the mNeonGreen signal colocalized with FM™ 4-64, a red fluorescent membrane dye, indicative of membrane localization for the fluorescently tagged DimA protein (Fig. 4d). Deletion of the N-terminal predicted signal peptide (*Δ*M1-D28) resulted in uniform cytoplasmic localization of mNeonGreen which highlights the importance of the N-terminal domain for the localization of DimA (Fig. 4d). Expression of the DimA^ΔM1-D28^ variant from an ectopic locus (*attB::P_dimA_-dimA^ΔM1-D28^*) in the *dimA* mutant failed to complement DimA function (Fig. S6). To further verify that DimA’s functional domain is localized in the periplasm, we generated a C-terminal fusion with alkaline phosphatase (PhoA) that is active only when it is in the periplasm ([36]). Both the WT and *ΔkinB* strains containing the DimA’-‘PhoA fusion were able to degrade p-nitrophenyl phosphate disodium salt hexahydrate (pNPP) substrate, showing activity of the PhoA enzyme and localization of the fusion in the periplasm (Fig. 4e). We infer that DimA functions in the periplasmic space.

**Fig. 4:**
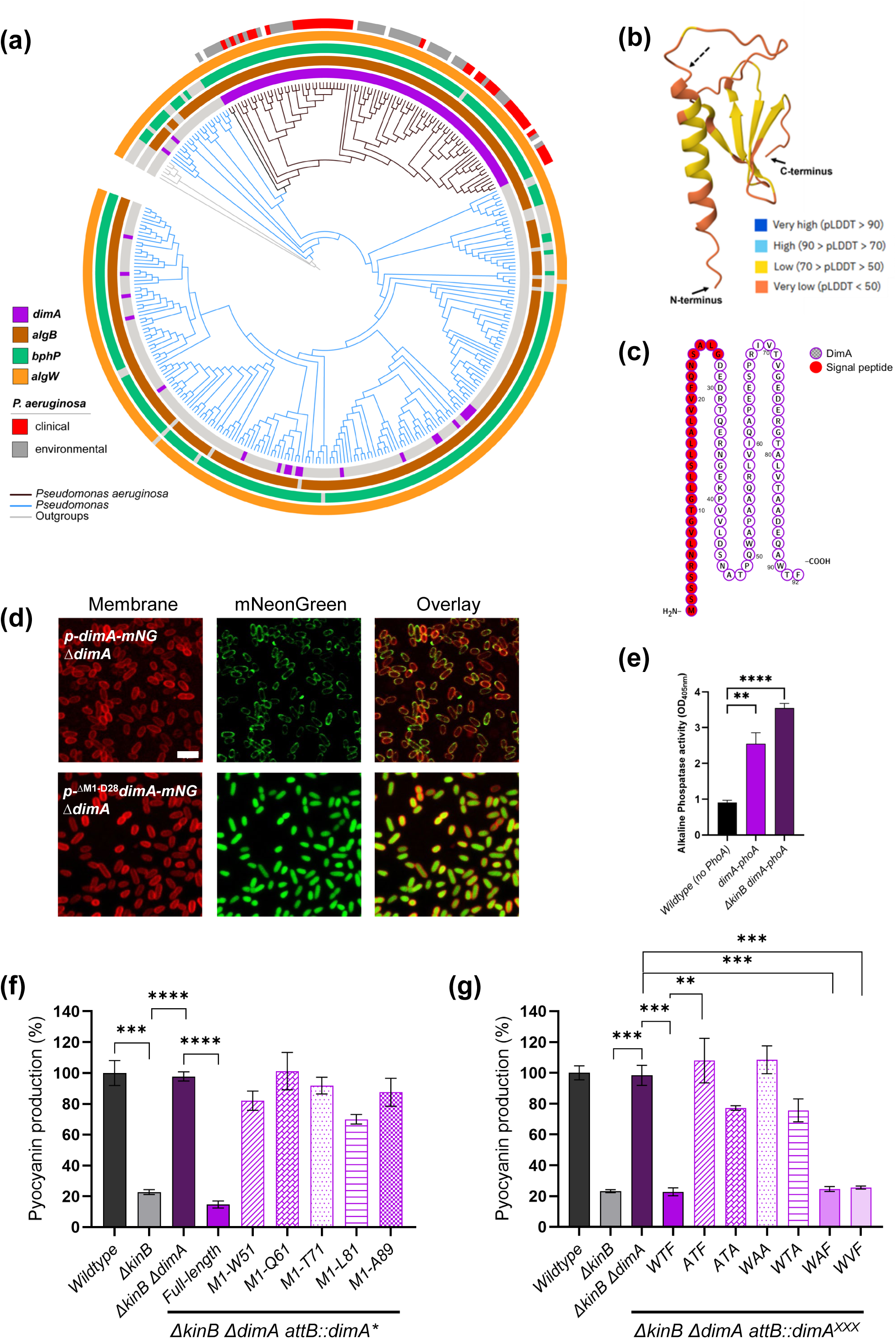
The C-terminal WTF motif of DimA is required for its function. (a) Phylogenetic distribution and presence/absence of select genes across *Pseudomonas* spp. Rings surrounding the tree depict metadata and gene presence/absence across genomes - 4 innermost rings: presence (colored) or absence (gray) of four regulatory genes—dimA (purple), algB (brown), bphP (teal), and algW (orange)—as determined by reciprocal BLAST against the P. aeruginosa PA14 genome. Outer ring: classification of *P. aeruginosa* strains as clinical (red) or environmental (gray). Colored branches in tree represent *P. aeruginosa* (black), other Pseudomonas spp. (blue), and outgroup genomes (gray). (b) AlphaFold structure prediction for DimA (PA14_20480) protein. The dashed arrow points at the predicted signal peptidase cleavage site. The N- and C-termini are shown by their respective arrows. The structure is colored according to the per-residue model confidence score (pLDDT). (c) Protter visualization plot of DimA amino acid sequence. The predicted signal peptide is colored red and the rest of the sequence is outlined in purple. (d) Confocal microscopy analysis of DimA and its N-terminal deletion mutant in *P. aeruginosa*. The cells were grown overnight in LB medium supplemented with 100μg/ml tetracycline and 1mM IPTG. Top row: PA14 *ΔdimA* strain harboring the plasmid pME6032-*dimA-mNeonGreen*, bottom row: PA14 *ΔdimA* strain harboring the plasmid pME6032-*dimA^ΔM1-D28^*-mNeonGreen fusion under induced conditions. Left to right: Membrane stained with FM^TM^4-64 (red), DimA-mNeonGreen fusion protein (green), and an overlay of both channels. Scale bar is 3 μm for all images. (e) Measurement of alkaline phosphatase activity. Overnight cultures of *P. aeruginosa* WT and *ΔkinB* strains bearing the *dimA-phoA* translational fusion were normalized for their growth at OD_600_=1. The phosphatase activity was measured at OD_405_ by the production of a yellow-colored product of the pNPP substrate. Error bars represent SEM of three biological replicates. Statistical significance was determined using t-test pairwise comparisons in GraphPad Prism software. ** P <0.005, **** P <0.0001. (f) Different truncation mutants of DimA were assayed for pyocyanin production. (g) Alanine substitutions in the WTF motif of DimA were tested for pyocyanin production. (f-g) Pyocyanin production (OD_695_) was measured and normalized to growth (OD_600_) in WT PA14 and the designated mutants grown overnight under ambient light conditions. Pyocyanin levels in WT PA14 were set to 100%. Error bars represent SEM of three biological replicates. Only pairwise comparisons that had p value <0.05 are denoted. Statistical significance was determined using t-test pairwise comparisons in GraphPad Prism software. **** P<0.0001, *** P<0.001, ** P <0.005

Next, to investigate how a periplasmic microprotein could affect the protein levels of the cytoplasmic response regulator AlgB, we generated several truncated versions of *dimA* and expressed them from its native promoter at an ectopic locus (*attB::P_dimA_-dimA^M1-W51^/dimA^M1-Q61^/dimA^M1-T71^/dimA^M1-L81^/dimA^M1-A89^/dimA^M1-F92^*). We performed pyocyanin production assay for these DimA variants in the sensitized *ΔkinB* mutant background and found that only the full-length DimA protein was able to repress pyocyanin production and even deletion of the last three amino acids (DimA_1-89_ or *ΔWTF*) completely abolished DimA’s function (Fig. 4f). To define the amino acid specificity of the WTF motif, alanine substitutions were made in the WTF motif and pyocyanin production was measured again as a proxy for the protein’s functionality. Interestingly, the alanine substitutions showed that pyocyanin production was repressed only when Trp90 and Phe92 were kept intact while the Thr91 was dispensable for the function of DimA (Fig. 4g). Conclusively, these data support that the C-terminal aromatic amino acids Trp90 and Phe92 are necessary for DimA function.

### DimA activates the RIP site-I protease AlgW

One way in which DimA might promote AlgB protein levels is via the RIP cascade that governs the availability of the alternate sigma factor AlgU. Furthermore, *algW* was hit in the same transposon screen that identified DimA (Fig. 1d, S1, S2). To test the idea that DimA could trigger AlgW mediated proteolysis of MucA, we first generated a markerless deletion of *algW* in the WT and *ΔkinB* mutant backgrounds and examined whether the absence of AlgW showed similar physiological responses as the deletion of *dimA*. Indeed, repression of pyocyanin production (Fig. 5a) and biofilm formation (Fig. 5b) under ambient light were abolished in the *ΔkinBΔalgW* mutant, mirroring the effects observed in the *ΔkinBΔdimA* mutant. These findings are consistent with AlgW being a critical interacting partner of DimA, mediating light-dependent repression of pyocyanin production and biofilm formation in *P. aeruginosa*. Next, to test our hypothesis further, we employed AlphaFold3 model prediction and found that DimA is predicted to bind through the WTF motif to the PDZ domain of AlgW, likely leading to catalytic activation of the protease domain (Fig. 5c, S7-8).

**Fig. 5:**
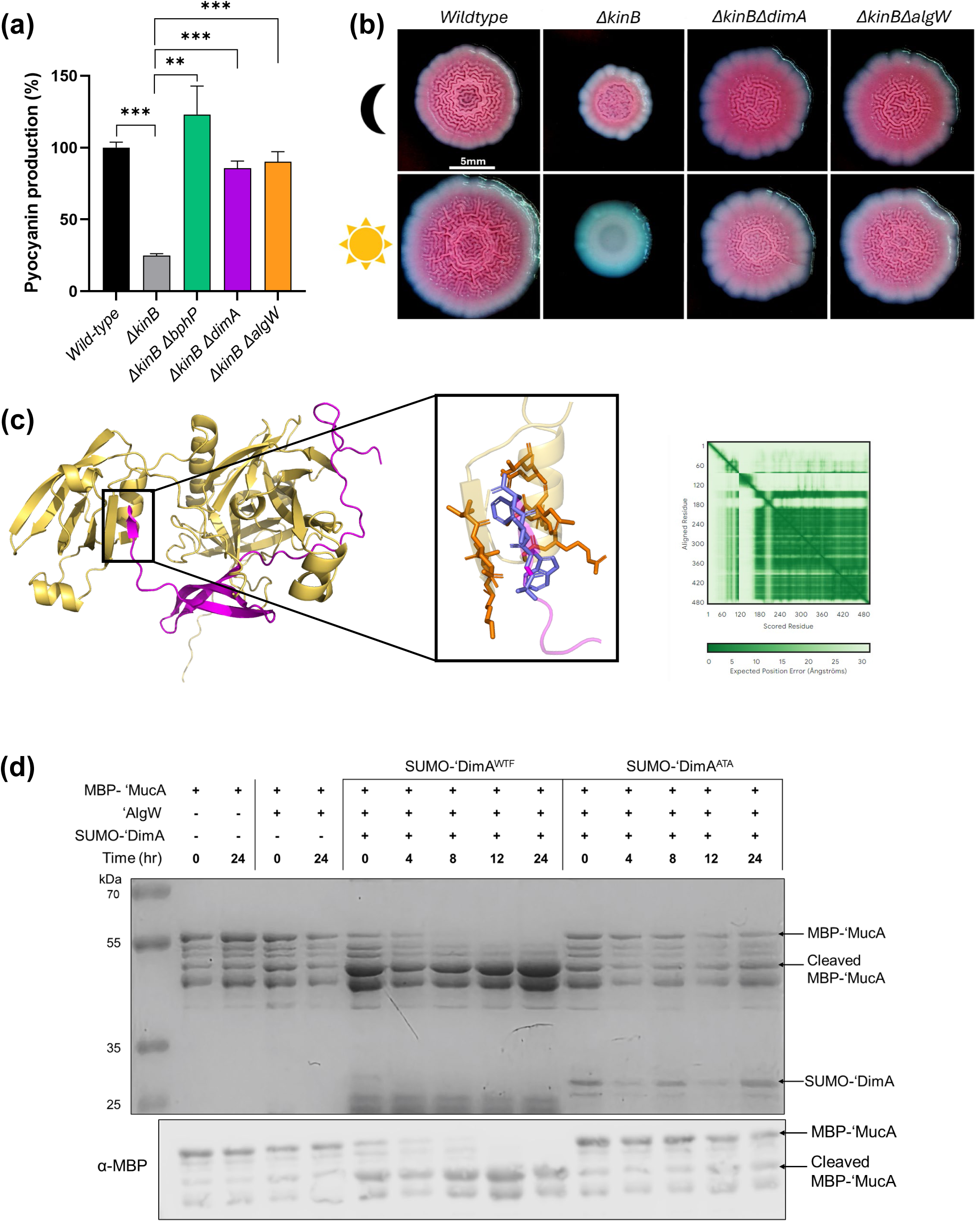
DimA is an activator of AlgW protease. (a) AlgW mutant was assayed for pyocyanin production. Pyocyanin production (OD_695_) was measured and normalized to growth (OD_600_) in WT PA14 and the designated mutants grown overnight under ambient light conditions. Pyocyanin levels in WT PA14 were set to 100%. Error bars represent SEM of three biological replicates. Only pairwise comparisons that had p value <0.05 are denoted. Statistical significance was determined using t-test pairwise comparisons in GraphPad Prism software. *** P<0.001 (b) Colony biofilm phenotypes of WT PA14 and the designated mutants on Congo red agar medium. The strains were grown for 72 h under dark (top row - denoted by the “black crescent” symbol) or ambient light conditions (bottom row – denoted by the “sun” symbol). Top-down view of stereoscope images. Scale bar is 5 mm for all images. Representative image from one of the triplicates is shown. (c) Alphafold3 multimer prediction model of AlgW periplasmic domain (colored in yellow) and DimA (colored in purple) structures (left). Confidence in inter- and intra-chain positioning is visualized using the Predicted Aligned Error (PAE) plot (right). In the PAE heatmap, each cell represents the expected positional error (in Ångströms), dark green indicates low expected positional error (high confidence), while light green to white indicates higher uncertainty. The interaction between the DimA WTF motif and AlgW PDZ domain has been boxed and the key residues involved in the interaction have been enlarged and displayed as sticks. (d) *In vitro* analysis of MBP-‘MucA proteolysis in the presence of AlgW protease and the DimA activator. SDS-PAGE gel stained with Coomassie Brilliant Blue (top panel) and corresponding anti-MBP immunoblot (bottom panel). Molecular weight markers (kDa) are shown on the left.

To determine if DimA activates AlgW directly, an *in vitro* proteolysis assay was conducted. Purified AlgW, MucA and DimA proteins were incubated together over a period of time and proteolysis of MucA was monitored via Coomassie Brilliant Blue staining as well as Western blot using MBP specific antibodies. Specifically, we generated an N-terminal fusion of maltose binding protein (MBP) with the periplasmic domain of MucA (MBP-‘MucA), an N-terminal fusion of 6xHis-SUMO tag to the periplasmic part (amino acids 29-92) of DimA (SUMO-‘DimA) and an N-terminal fusion of 6xHis tag to the periplasmic domain of AlgW (‘AlgW). Incubation of MucA with only AlgW did not result in any significant proteolysis of MucA (Fig. 5d). In contrast, the addition of DimA induced proteolysis of MucA as the levels of uncleaved MBP-‘MucA decreased over time while the levels of cleaved MucA increased concomitantly (Fig. 5d). To test the importance of the C-terminal WTF motif for activation of AlgW-mediated proteolysis, we purified a DimA variant where the WTF motif was changed to ATA (SUMO-‘DimA^ATA^). The purified DimA^ATA^ variant failed to activate AlgW-dependent degradation of MucA (Fig. 5d). Notably, in addition to the degradation of MucA we also observed degradation of DimA over time *in vitro*. However, heat-inactivated AlgW failed to degrade both MucA and DimA, showing that catalytically active AlgW is required for MucA and DimA proteolysis (Fig. S9). To confirm that AlgW is a specific protease, we incubated AlgW with bovine serum albumin (BSA) protein in the presence of DimA and did not observe any degradation of BSA over time (Fig. S9). Taken together, these data demonstrate that DimA directly activates AlgW protease function through its C-terminal WTF motif.

### DimA is a light-induced microprotein

The nucleotide sequences of the upstream promoter regions of *dimA* in *P. aeruginosa* strains are highly conserved (Fig. S10), suggesting that the regulation of its expression is universal across *P. aeruginosa* isolates. We found a putative AlgB binding motif ([37]) and hypothesized that *dimA* expression is induced by light via AlgB. To monitor the expression of *dimA*, we generated a chromosomal transcriptional reporter by integrating the β-galactosidase gene, *lacZ*, downstream of *dimA* in the PA14 genome. By measuring β-galactosidase activity, we had an estimate for transcriptional levels of *dimA*. Exposure to far-red light achieved striking upregulation of *dimA* in WT PA14 strain compared to the dark condition (Fig. 6a). Deletion of the response regulator *algB* or the photoreceptor *bphP* abrogated this induction, demonstrating their essential roles in *dimA* expression. Additionally, absence of the stress-induced alternative sigma factor *algU* did not affect the upregulation of *dimA* (Fig. 6a). These data establish that DimA is a photo-induced microprotein. Consistent with this, bioinformatic analysis of publicly available transcriptomes showed higher *dimA* expression in samples that are likely “light-exposed” (for example, human burn wound) compared to those that are likely “dark” (for example, pneumonia) (Fig. 6b, S11).

**Fig. 6:**
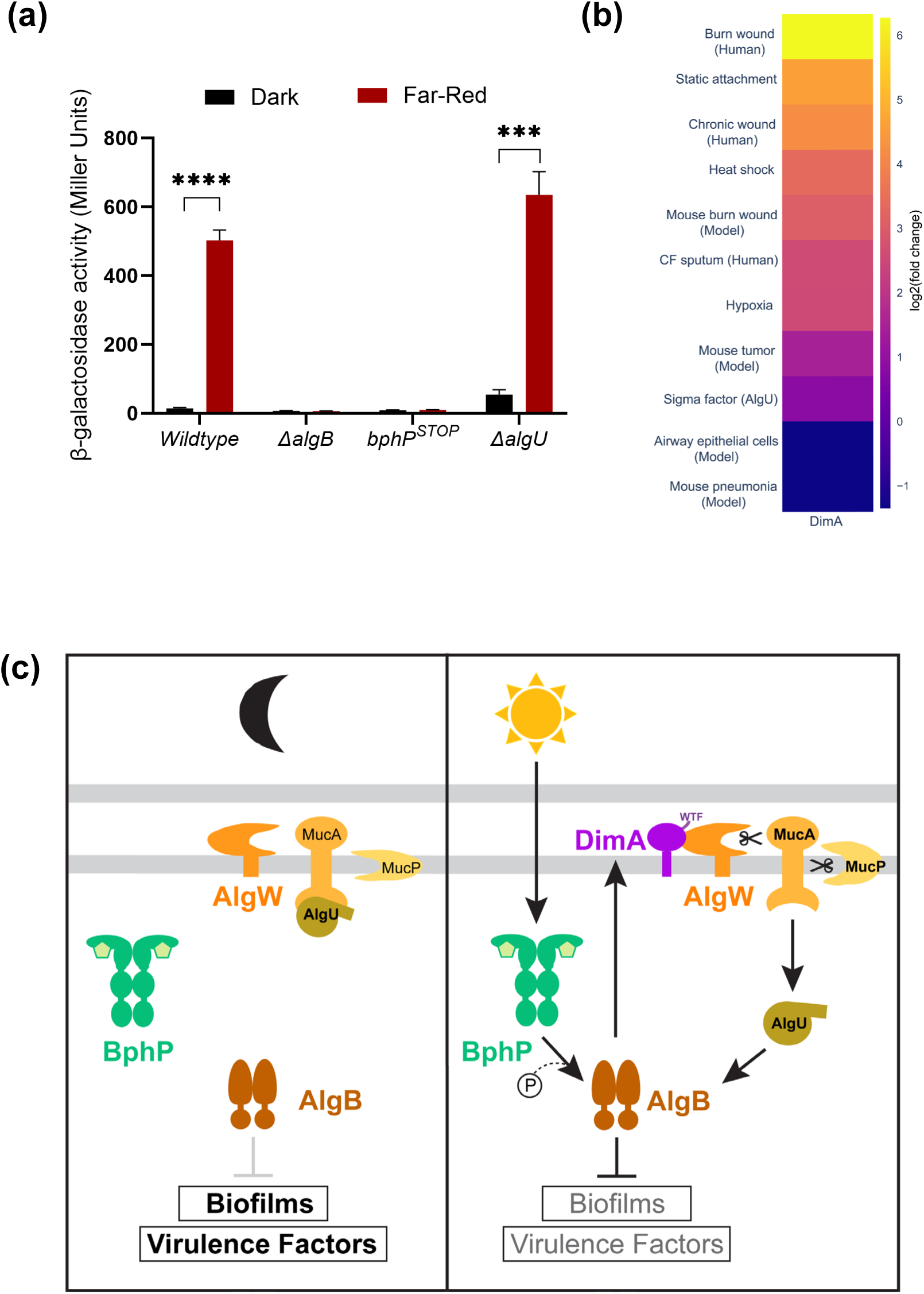
Light induces *dimA* expression in an AlgB-dependent manner. (a) β-galactosidase assays of *PdimA-lacZ* transcriptional fusion for background genotypes indicated on the X-axis and grown in dark and far-red light conditions. Error bars represent SEM of three biological replicates. Only pairwise comparisons that had *p* value <0.05 are denoted; statistical significance was determined using GraphPad Prism software. *** P <0.001, **** P <0.0001. (b) Heatmap showing the log₂ fold change in expression of the *dimA* gene across a subset of publicly available RNA-seq datasets, relative to LB-grown *P. aeruginosa* PA14. Extended analysis in S11. (c) Model for DimA-dependent activation of photo sensing. (Left) In the absence of light, BphP (green) is inactive, AlgB (brown) is not phosphorylated and photo-sensing signaling is in an “OFF” state. (Right) Light activates BphP-AlgB TCS, P∼AlgB in turn activates the expression of DimA (purple). DimA triggers regulated intramembrane proteolysis via the site-I protease AlgW (orange) to release the sigma factor AlgU (tan) that leads to increased AlgB levels which in turn upon phosphorylation by BphP represses biofilms and virulence factors in the photo sensing “ON” state. Arrow indicates activation and T-bar indicates inhibition.

## DISCUSSION

In this study, we have identified and characterized DimA, a novel 92-amino acid microprotein that functions as a master activator of the photo-sensing response in *P. aeruginosa*. Our findings reveal a sophisticated regulatory mechanism through which *P. aeruginosa* senses and responds to light via a complex signaling cascade involving DimA, the BphP-AlgB two-component system, and the regulated intramembrane proteolysis (RIP) pathway. DimA serves as a critical intermediary in this photo-sensing pathway, amplifying the light-induced signal through a positive feedback mechanism. Positive feedback in transcriptional regulation creates efficiency by limiting production of inactive regulatory proteins that require activation (such as phosphorylation) to function ([38], [39]). We propose that the DimA-mediated regulatory mechanism prevents wasteful synthesis of AlgB protein in the absence of light (Fig. 6c) and reduces the risk of inappropriate activation by non-physiological partners, as reported in other bacterial systems like sporulation in *Bacillus subtilis* and virulence pathways in *Salmonella enterica* ([40], [41]).

The molecular mechanism by which DimA regulates AlgB involves the RIP pathway. Our structural and functional analyses revealed that DimA is a membrane-localized microprotein with a periplasmic C-terminal domain containing a critical WTF motif. This motif bears striking similarity to the WVF motif in MucE, a known activator of the site-I protease AlgW which initiates the cleavage of MucA in the regulated intramembrane proteolysis pathway of *P. aeruginosa* ([42], [43]). Indeed, our AlphaFold3 modeling and experimental data strongly suggest that DimA interacts with AlgW through its WTF motif, activating the protease and triggering the RIP cascade that ultimately leads to the release of the alternative sigma factor AlgU and subsequent upregulation of *algB*. Therefore, our discovery that DimA activates the same proteolytic cascade but in response to light stimuli represents a novel integration of environmental sensing with this established RIP signaling pathway. Furthermore, our phylogenetic analyses identified DimA homologs in other Pseudomonads such as *P. chlororaphis*, *P. syringae*, and *P. arsenicoxydans*, and AlphaFold predictions suggest that these DimA homologs interact with the corresponding AlgW proteases (Fig. S4, S5, S8). We speculate that DimA plays a similar role in these bacteria that contain functional BphP-AlgB photo-sensing cascades ([18], [44]).

The identification of DimA adds to our growing understanding of the critical role of microproteins in bacterial signaling networks. Microproteins, typically encoded by small open reading frames (sORFs), have emerged as important regulatory elements in diverse cellular processes including ion transport, oxidative phosphorylation, and stress signaling ([45], [46]). These small proteins, often comprising the “dark matter” of bacterial proteomes, remain largely uncharacterized despite being widely distributed across all domains of life ([47], [48], [49], [50], [51]). Of the characterized microproteins, some function as allosteric regulators of larger proteins or protein complexes, modifying or fine-tuning their activities, while others serve as independent signaling molecules. Small proteins involved in the regulation of two-component systems include SafA and MgrB that function as negative regulators of the histidine kinase PhoQ in the PhoQ/PhoP two-component system in *Escherichia coli* and *Salmonella enterica*, respectively ([52], [53], [54], [55]). DimA exemplifies the paradigm of microproteins as critical regulatory elements in several ways. First, like many functional microproteins, DimA contains a transmembrane domain that likely anchors it to the bacterial membrane where it can effectively interface between different cellular compartments - in this case, between the periplasmic and cytoplasmic environments. Second, DimA functions not through enzymatic activity of its own, but as an allosteric regulator by modulating the activity of a larger protein complex (the AlgW protease) through a specific protein-protein interaction via its C-terminal WTF motif. Third, DimA operates as a signal amplifier as it enhances an initial environmental signal (light) into a robust cellular response (changes in virulence and biofilm formation). In essence, DimA represents a perfect example of how microproteins, despite their small size, can serve as sophisticated molecular switches in complex bacterial signaling networks, connecting environmental sensing to coordinated cellular responses.

The high conservation of DimA across both clinical and environmental *P. aeruginosa* isolates underscores its fundamental role in *P. aeruginosa* physiology and environmental adaptation. As an environmental cue, light provides bacteria with critical information about their location and recent studies have shown that growth in darkness promotes virulence-associated traits and enhanced pathogenic potential in *P. aeruginsa* ([56], [57]). From a clinical perspective, these findings open promising therapeutic avenues against P. aeruginosa, a significant opportunistic pathogen associated with serious infections in immuno-compromised individuals and cystic fibrosis patients. Indeed, artificial overexpression of DimA homolog in the *P. aeruginosa* clinical isolate IHMA879472 led to increased resistance to human plasma ([58]). Specifically, targeting the DimA-mediated photo-sensing pathway could offer novel strategies to disrupt biofilm formation and virulence factor production, two critical determinants of *P. aeruginosa* pathogenicity in clinical settings. This approach aligns with emerging interest in developing anti-virulence therapies that do not directly kill bacteria but rather interfere with their pathogenic capabilities, potentially reducing selective pressure for resistance development.

In sum, our study identifies DimA as a critical regulator of the photo-sensing response in *P. aeruginosa* and elucidates its mechanism of action through activation of the RIP pathway. This work opens several promising avenues for future research in this important pathogen. First, kinetic and structural studies to elucidate the precise molecular interactions between DimA and the periplasmic domain of AlgW should provide insights into the proteolysis of DimA, potentially guiding the development of peptide inhibitors or small molecules that could modulate this interaction. Second, exploring the potential crosstalk between the photo-sensing pathway and other environmental stress responses, such as oxidative stress or antibiotic exposure, could reveal integrated regulatory networks. Finally, since *P. aeruginosa* infections often involve polymicrobial communities, examining how DimA-mediated photo-sensing affects interspecies interactions could yield insights into complex microbial behaviors in clinical and environmental settings.

## MATERIALS AND METHODS

### Strains and growth conditions

*P. aeruginosa* UCBPP-PA14 strain was grown in lysogeny broth (LB) (10 g tryptone, 5 g yeast extract, 5 g NaCl per L) and on LB plates fortified with 1.5% agar at 37°C. When appropriate, antimicrobials were included at the following concentrations: 400 µg/mL carbenicillin, 50 µg/mL gentamycin, 100 μg/mL irgasan, 500μg/mL trimethoprim, 50 µg/mL kanamycin, and 100 μg/mL tetracycline.

### Strain construction

Strains and plasmids were constructed as described previously ([59]). To construct marker-less in-frame chromosomal deletions in *P. aeruginosa*, DNA fragments flanking the gene of interest were amplified, assembled by the Gibson method, and cloned into pEXG2 ([60]). The resulting plasmids were used to transform *Escherichia coli* SM10λ*pir*, and subsequently, mobilized into *P. aeruginosa* PA14 via biparental mating. Exconjugants were selected on LB containing gentamicin and irgasan, followed by recovery of deletion mutants on LB medium containing 5% sucrose. Candidate mutants were confirmed by PCR. The complementation strain for PA14_20480 and its variants were cloned into pUC18T-mini-Tn*7*T-Tp plasmid ([61]). The resulting plasmids were transformed to *Escherichia coli* DH5α, and mobilized into *P. aeruginosa* PA14 via quad-parental mating using as helper strains DH5α/pTNS3 and DH5α/pRK2013 to facilitate Tn7 transposition at the *attB* site as described here ([61]). Insertion of the Tn7 transposon region was confirmed by PCR followed by Sanger sequencing for verifying the truncated versions and point mutations of *PA14_20480*. Similarly, the construction of the luminescent *lecA* reporter strains was performed by quad-parental mating between PA14 strains, DH5α/pUC18T-mini-Tn*7*T-Tp-*PlecA-luxCDABE,* and the helper strains DH5α/pTNS3 and DH5α/pRK2013.

To construct the transcriptional reporter fusions of P*20480ΩlacZ*, ∼1000 bp of DNA upstream of the stop codon of *PA14_20480* and the DNA encoding the *lacZ* open-reading frame were amplified using *P. aeruginosa* PA14 genomic DNA and the plasmid pIT2 as templates, respectively. Next, a DNA fragment of ∼1000 bp downstream of *PA14_20480* was amplified from *P. aeruginosa* PA14 genomic DNA. The three DNA fragments were assembled by the Gibson method and cloned into pEXG2. The resulting plasmid was used to transform *E. coli* SM10λ*pir*, and subsequently mobilized into *P. aeruginosa* PA14 WT and the Δ*algB*, Δ*algU, bphP*STOP mutants via biparental mating as described above.

To construct the translational fusion of PA14_20480-PhoA, ∼1000 bp of DNA upstream and downstream of *PA14_20480,* the DNA encoding the *phoA* open-reading frame, were amplified using *P. aeruginosa* PA14 genomic DNA and the plasmid pCM639 as templates, respectively ([62]). The three DNA fragments were assembled by the Gibson method and cloned into pEXG2. The resulting plasmid was used to transform *E. coli* SM10λ*pir*, and subsequently mobilized into *P. aeruginosa* PA14 WT.

For the construction of PA14_20480-mNeonGreen or PA14_20480_Cterm_-mNeonGreen the open-reading frame of PA14_20480 or PA14_20480*Δaa2-28*, and the *mNeonGreen* were amplified using *P. aeruginosa* PA14 genomic DNA and the plasmid p*mNeonGreen*-N1 as templates, respectively ([63]). The two DNA fragments were assembled by Gibson method and cloned into pME6032 plasmid ([64]).

Protein production constructs were generated by amplifying the *algW^Δaa1-32^, mucA^Δaa1-105^, dimA^Δaa1-28^* coding regions and cloning them in pET28b (Addgene, Watertown, MA, USA), pMAL-c6T (New England Biolabs, Ipswich, MA, USA), and pTB146 ([65]) expression vectors to obtain pET28b-His6-‘AlgW, pMAL-MBP-‘MucA, and pTB146-SUMO-‘DimA, respectively.

### Transposon mutagenesis screen

We generated a sensitized reporter strain by deleting the genes encoding the known negative regulator of photo sensing, KinB and the biliverdin producing heme oxygenases BphO and HemO ([66], [67]). This sensitized Δ*kinB*Δ*bphO*Δ*hemO attB::PlecA’-‘luxCDABE* strain was subjected to Mariner-based transposon mutagenesis using pBT20 vector ([68]). Our rationale was that inactivation of a gene(s) encoding a component that promotes photo sensing would sever the connection between light and repression of the *lecA-lux* reporter. Insertion mutants were selected on LB agar containing 100 µg/mL gentamycin and 100 μg/mL irgasan was included in the agar to counter select against the *E. coli* donor. The plates were incubated at 25°C for 48 hours in the presence of light and screened for colonies expressing increased luminescence as detected by an Amersham ImageQuant 800 (Cytiva) imager. Transposon insertion locations were determined by arbitrary PCR and sequencing as described previously ([69]).

### Luminescence reporter assay

PA14 strains harboring the chromosomally encoded *attB::*P*lecA-luxCDABE* were grown overnight at 37°C in LB growth medium. The next day, 5μL of the overnight cultures were used to inoculate 1mL cultures and grew them at 37°C in LB medium until they reached OD_600_=0.5. 10-fold serial dilutions were carried out and plated on LB agar plates. The plates were incubated in different light conditions for 48 h and then measured the chemiluminescence with 3 min exposure time using an Amersham ImageQuant 800 (Cytiva) imager. Quantification of the luminescent values was performed with Image Lab 6.1 software (Bio-Rad). The luminescent values were normalized to the culture growth from the CFU measurement of the serial dilutions.

### Colony biofilm assay

One microliter of *P. aeruginosa* cultures (OD=0.5) grown at 37°C in LB broth was spotted onto 60 x 15 mm Petri plates containing 10 mL 1% Tryptone medium supplemented with 40 mg/L Congo red, 20 mg/L Coomassie brilliant blue dyes, and solidified with 1% agar. Colonies were grown at 26°C under dark, ambient light, or Far-Red light (730nm, 1 Watt/m^2^) conditions, and images were acquired after 120 h using a Zeiss AxioZoom v16 microscope.

### Pyocyanin Assay

PA14 strains were grown overnight in LB liquid medium at 37 °C with shaking at 220 rotations per minute (rpm). The cells were pelleted by centrifugation at 15,000 × *g* for 2 min, and the clarified supernatants were passed through 0.22-μm filters (Millipore, Burlington, MA, USA) into clear plastic cuvettes. The OD_695_ of each sample was measured on a spectrophotometer (Genesys 20, ThermoScientific) and normalized to the culture cell density, which was determined by OD_600_.

### Protein Overexpression and Purification

#### His6-AlgB

The pET28b-His6-AlgB protein production and purification for the production of the anti-AlgB antibody followed the process described here ([18]).

#### His6-‘AlgW

The pET28b-His6-AlgW vector was transformed into BL21(DE3) and the culture was grown approximately to 0.8 OD_600_ in 1 L of LB supplemented with 50 μg/mL kanamycin at 37 °C with shaking at 220 rpm. Protein production was induced by the addition of 1 mM IPTG and incubation of the culture overnight at 37 °C was followed. The cells were pelleted by centrifugation at 4000rpm for 20 min and resuspended in AlgW-lysis buffer (25 mM Tris-HCl (pH 7.5), 150 mM NaCl, and 5% glycerol). The cells were lysed by sonication (20s pulses for 30min). The lysate was centrifuged at 12100rpm for 30 min at 4 °C. The resulting clarified supernatant was combined with Ni-NTA resin (Novagen) and incubated for 1 h at 4 °C. The bead/lysate mixture was loaded onto a 1.5-cm Econo-Column® (Bio-Rad, Hercules, CA, USA), the resin was allowed to pack, and then it was washed with AlgW-wash buffer (25 mM Tris-HCl (pH 7.5), 150 mM NaCl, and 5% glycerol, 25 mM imidazole) Resin-bound His6-AlgW was eluted twice with 3 column volumes (CV) AlgW-lysis buffer containing 300 mM imidazole. Fractions were analyzed by SDS-PAGE, and the gel was stained with Coomassie brilliant blue to assess His6-AlgW purity. Purified protein was dialyzed in degradation buffer (50 mM sodium phosphate (pH 7.4), 200 mM KCl, and 10% glycerol) and stored at −80 °C.

#### His6-SUMO-‘DimA and His6-SUMO-‘DimA_ATA_

The pTB146-SUMO-‘DimA and pTB146-*SUMO-‘DimA_ATA_* vector were transformed into BL21(DE3) and the culture was grown approximately to 0.6 OD_600_ in 1 L of LB supplemented with 200 μg/mL carbenicillin at 37 °C with shaking at 220 rpm. Protein production was induced by the addition of 1 mM IPTG and incubation of the culture overnight at 37 °C was followed. The cells were pelleted by centrifugation at 4000rpm for 20 min and resuspended in DimA-lysis buffer (50 mM Na_2_HPO_4_, 300mM NaCl and 10mM imidazole, 1 tablet of cOmplete™, Mini Protease Inhibitor Cocktail - Sigma-Aldrich). The cells were lysed by sonication (20s pulses for 30min). The lysate was centrifuged at 12100rpm for 30 min at 4 °C. The resulting clarified supernatant was combined with Ni-NTA resin (Novagen) and incubated for 1 h at 4 °C. The bead/lysate mixture was loaded onto a 1.5-cm Econo-Column® (Bio-Rad, Hercules, CA, USA), the resin was allowed to pack, and then it was washed with DimA-wash buffer (50 mM Na_2_HPO4, 300mM NaCl and 30mM imidazole). Resin-bound *His6-SUMO-‘DimA* was eluted with 2 CV fractions of DimA-lysis buffer containing 50mM, 100mM, 250mM, and 500mM imidazole. Fractions were analyzed by SDS-PAGE, and the gel was stained with Coomassie brilliant blue to assess SUMO-DimA purity. Purified protein was dialyzed in degradation buffer and stored at −80°C.

#### MBP-‘MucA

The pMAL-MBP-‘MucA vector was transformed into BL21(DE3) and the culture was grown approximately to 0.5 OD_600_ in 1 L of LB supplemented with 200 μg/mL carbenicillin and 0.2% glucose at 37°C with shaking at 220 rpm. Protein production was induced by the addition of 1 mM IPTG and incubation of the culture overnight at 16°C was followed. The cells were pelleted by centrifugation at 4000rpm for 20 min and resuspended in MBP-‘MucA lysis buffer (20mM Tris-HCl, 200mM NaCl, 1mM EDTA, and 1 tablet of cOmplete™, Mini Protease Inhibitor Cocktail). The cells were lysed by sonication (20s pulses for 30min). The lysate was centrifuged at 12100rpm for 30 min at 4°C. The resulting clarified supernatant was combined with amylose resin (New England Biolabs Inc.) and incubated for 1 h at 4°C. The bead/lysate mixture was diluted 1:5 in MBP-‘MucA lysis buffer loaded onto a 1.5-cm Econo-Column® (Bio-Rad, Hercules, CA, USA), the resin was allowed to pack. Then, it was washed with 12CV of MBP-‘MucA lysis buffer and eluted in 1CV of MBP-‘MucA lysis buffer containing 10mM maltose. Fractions were analyzed by SDS-PAGE, and the gel was stained with Coomassie brilliant blue to assess MBP-MucA purity. Purified protein was dialyzed in degradation buffer and stored at −80°C.

### Western Blot Assays

PA14 strains were grown overnight at 37°C in LB medium. 1 ml culture of cells was harvested and resuspended in 200μl of 2× SDS-PAGE Laemmli (Bio-Rad Laboratories, Inc., Hercules, CA, USA). The samples were heated for 10 minutes at 95°C. 10 μL samples were loaded onto 10% SDS-PAGE gels and were subjected to electrophoresis. Proteins were transferred to nitrocellulose membranes and afterwards blocked with 5% skim milk in TBS at room temperature for 1 h. Following, an incubation with primary antibodies anti-FLAG (Sigma-Aldrich), anti-AlgB (house-made by Cocalico Biologicals Inc., Denver, PA, USA), anti-RNAP (Abcam, Inc., Cambridge, UK), and anti-HA (R&D Systems) at 1:5000 dilution in 5% skim milk in TBS was performed overnight at 4°C on a rocking platform. Membranes were washed 3 times with TBS-Tween 20 at room temperature for 10 min on a rocking platform. Incubation with the secondary antibody anti-rabbit IgG-Peroxidase produced in goat (Sigma-Aldrich) was followed for 1 h at room temperature. The membranes were washed again for 3 times with TBS-Tween 20 for 10 min and subsequently developed with a SuperSignal West Femto Kit (Thermo Scientific) and captured with an Amersham ImageQuant 800 (Cytiva) imager. Quantification of the western blots was performed with Image Lab 6.1 software (Bio-Rad).

### Fluorescence Microscopy

PA14 strains bearing the pME6032-PA14_20480-mNeonGreen or the pME6032-PA14_20480_Cterm_-mNeonGreen plasmid were grown overnight at 37°C in LB medium containing 100μg/ml tetracycline and 1mM IPTG. 1 ml of culture was pelleted and washed once with 1ml PBS. The pellets were resuspended in 50μl solution of FM^TM^4-64 dye (Invitrogen) dissolved in PBS at 10μg/ml and incubated in dark for 1 min. The samples were washed once with 1 ml PBS and then resuspended in 100μl PBS final volume. The samples were loaded onto a microscope slide and covered with 1mm poly-L-Lysine coated coverslip. Micrographs were acquired at 60x magnification with immersion oil in a Stellaris 5 (Leica) microscope. Image processing was performed in Fiji platform (Fiji-2012).

### Alkaline phosphatase Assay

PA14 strains with the *PA14_20480-phoA* translation fusion were grown overnight at 37°C in LB medium. The cultures were diluted to OD_600_=1 and 1 ml of culture was pelleted by centrifugation at 15,000 x g for 1 min. The pellets were washed once with 1ml Tris pH=8 and then resuspended in 200ul of substrate p-nitrophenyl phosphate disodium salt hexahydrate (pNPP) at 1mg/ml (Sigma). The substrate was resuspended in 0.1 M glycine buffer (pH 10.4), with 1 mM MgCl2 and 1 mM ZnCl2. The samples were incubated in the dark at room temperature for 10 min. To stop the reaction 50ul of 3M NaOH were added. The optical density at 405nm was measured using a Synergy Neo2 microplate reader (Agilent BioTek).

### .β-galactosidase assay

PA14 strains with the *PA14_20480ΩlacZ* transcriptional fusion were grown overnight at 37°C in LB medium and 1mL cultures were collected pelleted. The pellets were resuspended in 1mL Z-buffer with the addition of 200μg lysozyme for permeabilization. Samples were then incubated at 30°C for 15 min and diluted 1:2 in a total volume of 500μL. The colorimetric reaction initiated with addition of 100μl of 4mg/mL ONPG (ortho-Nitrophenyl-β-galactoside) dissolved in Z-Buffer to the samples and time was recorded. The samples were incubated at 30°C until sufficient color change was observed. The reaction was quenched by 250μL 1M Na_2_CO_3_. The optical density of each sample was then measured at 420nm and 550nm. Standard activity was calculated in Miller units by this formula: ((OD_420_ – (OD_550_*1.75))*1000)/ (time * volume lysate * OD_600_).

### MucA proteolysis assay

All the added proteins had been dialyzed in the same degradation buffer (50 mM sodium phosphate (pH 7.4), 200 mM KCl, and 10% glycerol). 5uM of MBP-‘MucA, 1uM of ‘AlgW, and 5uM of SUMO-‘DimA or SUMO-‘DimA_ATA_ were mixed, and the reaction was incubated at 37°C. Samples were collected after stopping the reactions by adding 4x Laemmli Sample Buffer (Bio-Rad Inc.) containing 100mM DTT and heating the samples at 95°C for 10min. For heat inactivation of ‘AlgW and SUMO-‘DimA the proteins were heated at 95°C for 10min and then added to the mixture.

The samples were run in SDS-PAGE, and the gel was stained with Coomassie brilliant blue to assess MBP-‘MucA and SUMO-‘DimA cleavage. MBP-‘MucA cleavage was also examined by performing Western blot analysis using α-MBP Peroxidase conjugated antibody (Sigma-Aldrich) at a 1:7500 dilution and following the Western blot procedure described above.

## ACKNOWLEDGEMENTS

We thank the DFI Microbial Metagenomics Facility for help with RNA sequencing. We thank all members of the Mukherjee lab for thoughtful discussions. Research reported in this publication was supported by the National Institute of General Medical Sciences of the National Institutes of Health (NIH) under Grants R35GM150803 and R00GM129424 and the Searle Scholars Program Grant SSP-2022-104 to S.M. We are grateful to Margot and Robert Haselkorn and Virginia Aronson for their generous gift of equipment. The content of this study is solely the responsibility of the authors and does not necessarily represent the official views of the funding agencies. The funders had no role in study design, data collection and analysis, decision to publish, or preparation of the manuscript.

## CONFLICT OF INTEREST

The authors declare that they have no conflict of interest.

## DATA AVAILBILITY

All relevant data are provided within the manuscript and the supplementary information.

## SUPPLEMENTAL FIGURES

**Supplemental Figure 1:**
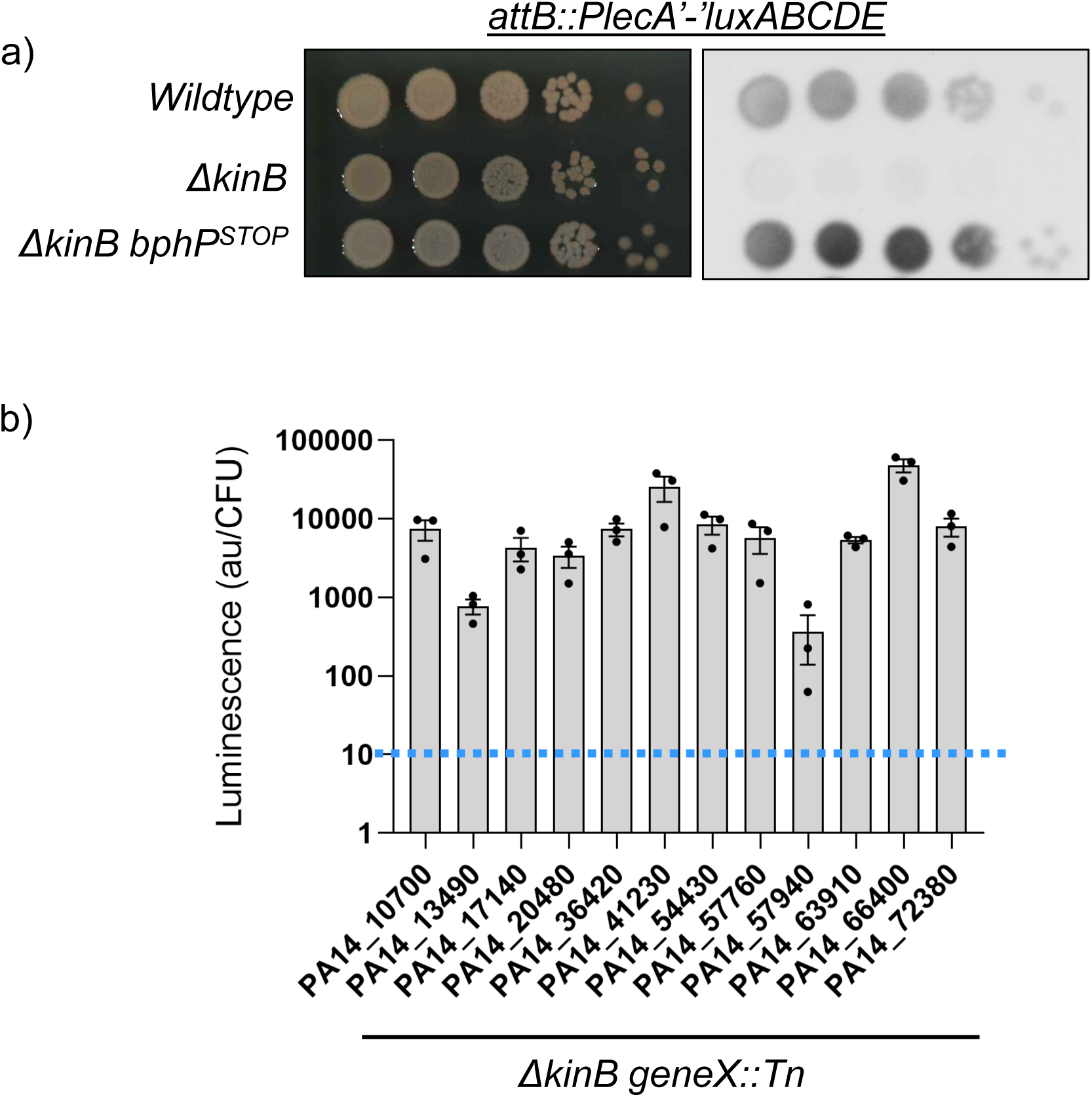
A *lecA-lux* based screen for positive regulators of photo sensing. a) Images of serial plating of designated strains depicting growth and *lecA-lux* expression. b) Quantification of *lecA-lux* activity for Tn5 candidates. Blue dotted line indicates activity from a parent *ΔkinB* mutant strain under ambient light conditions.

**Supplemental Figure 2:**
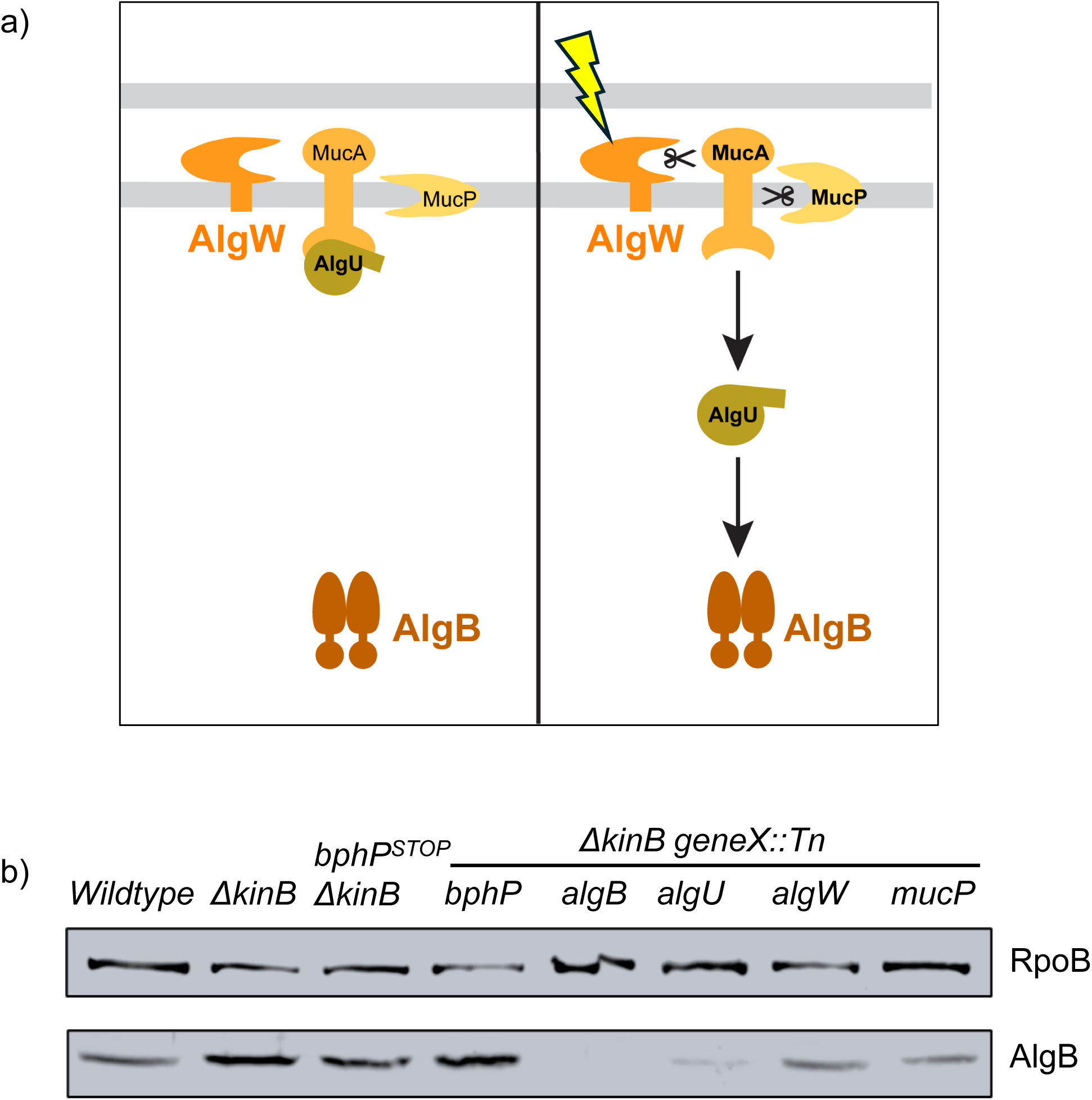
RIP mutants have low AlgB levels. a) Schematic of the RIP pathway. Outer membrane stress activates the site-I protease AlgW (orange) to cleave the anti-sigma factor MucA (mustard) which is subsequently cleaved by the site-II protease MucP (yellow) to release the sigma factor AlgU (tan) that leads to increased AlgB levels. Arrow indicates activation and T-bar indicates inhibition. b) Western blot analysis of AlgB protein levels using custom-raised antibodies. Overnight cultures of *P. aeruginosa* wildtype and the designated mutant strains were grown in the presence of ambient light. RpoB was used as a loading control.

**Supplemental Figure 3:**
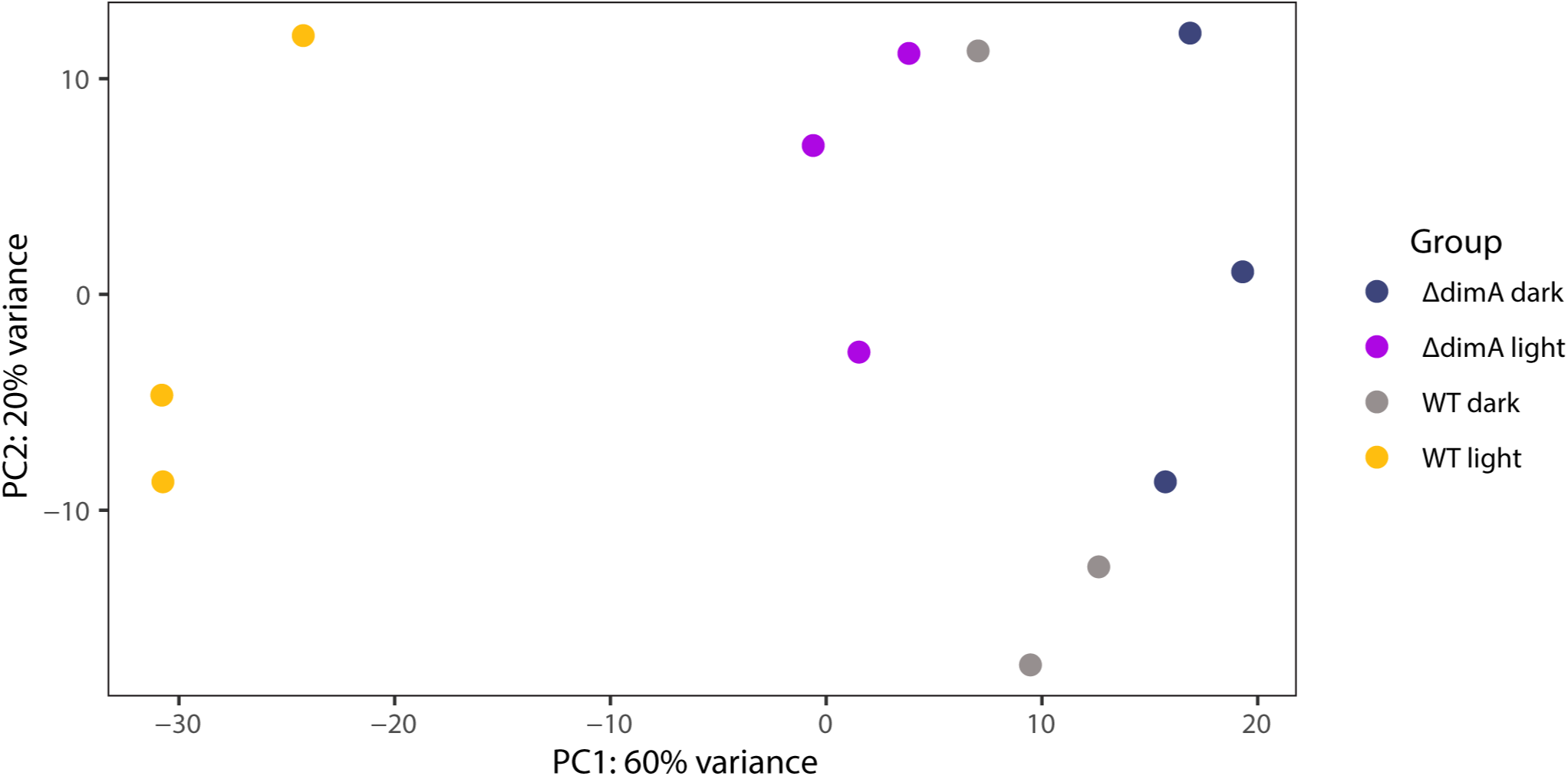
Principal component analysis (PCA) for RNA-seq on samples from the indicated strains.

**Supplemental Figure 4:**
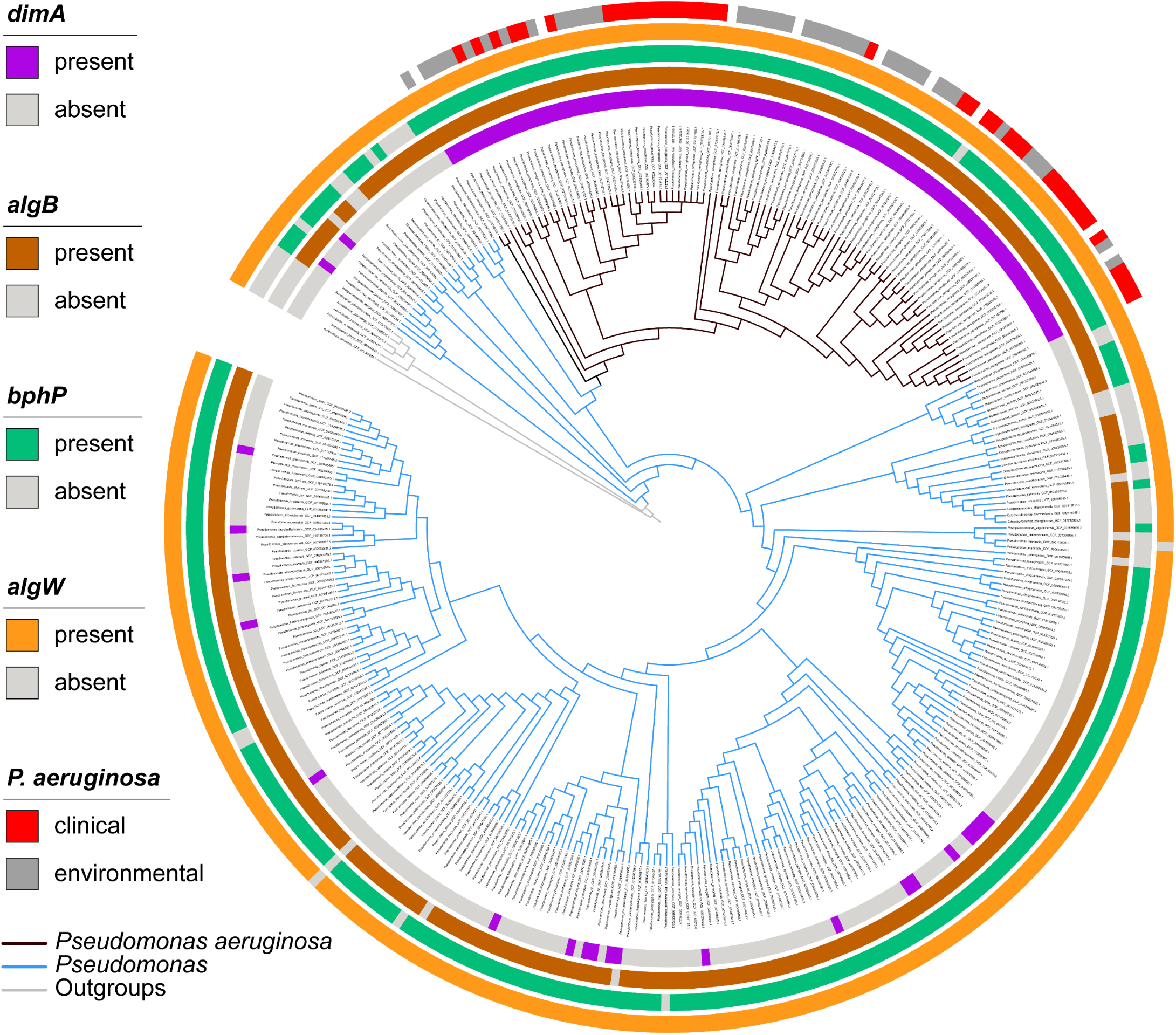
Phylogenetic tree from figure 4 reproduced at higher resolution.

**Supplemental Figure 5:**
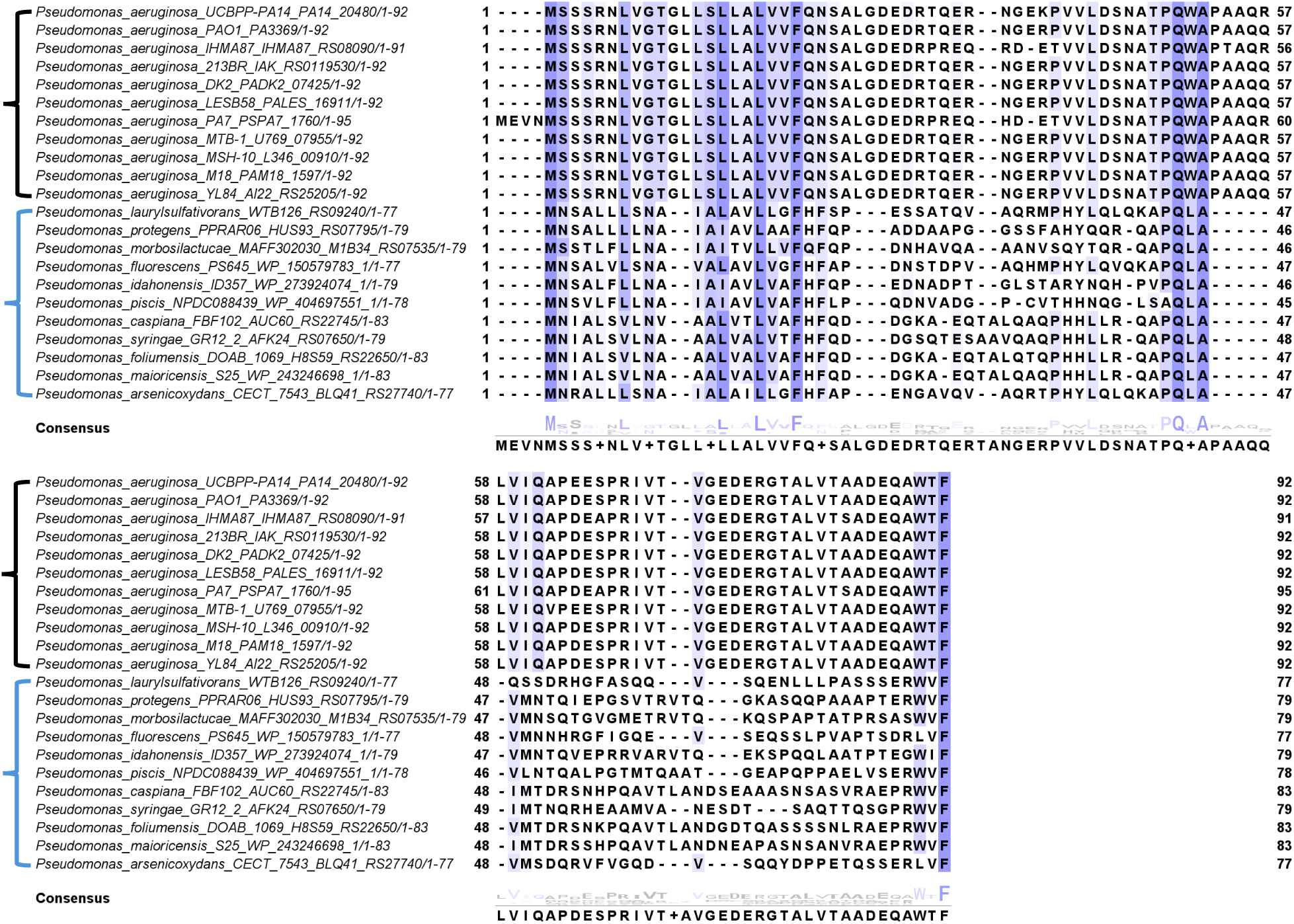
Multiple sequence alignment (MSA) for DimA homologs. Primary sequence alignment of DimA from representative *Pseudomonas aeruginosa* isolates (black cluster) and other *Pseudomonas spp.* (blue cluster). Conserved residues are highlighted in purple based on a ≥30% identity threshold, with increased color intensity indicating higher conservation.

**Supplemental Figure 6:**
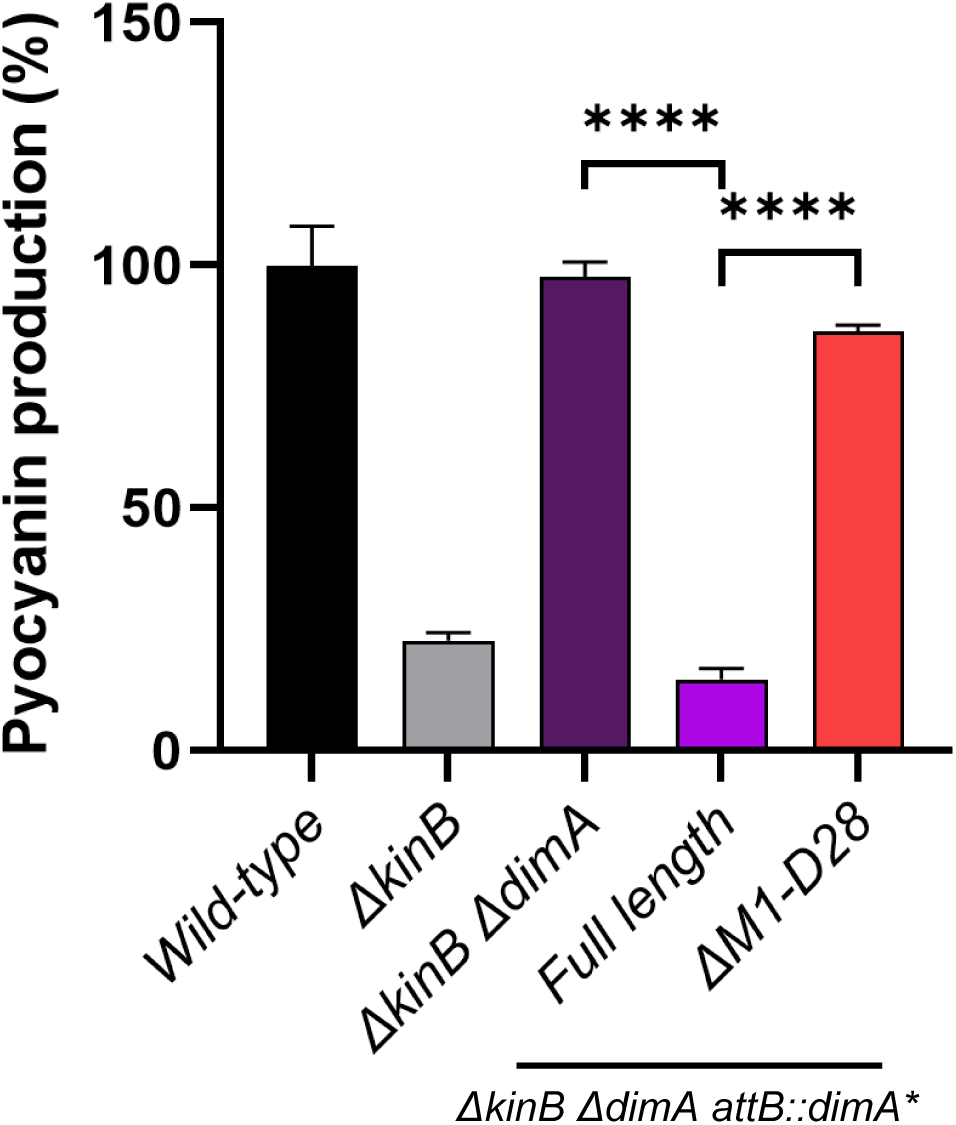
N-terminal signal peptide is required for DimA function. Pyocyanin production (OD_695_) was measured and normalized to growth (OD_600_) in WT PA14 and the designated mutants grown overnight under ambient light conditions. Pyocyanin levels in WT PA14 were set to 100%. Error bars represent SEM of three biological replicates. Statistical significance was determined using t-test pairwise comparisons in GraphPad Prism software. **** P <0.0001.

**Supplemental Figure 7.**
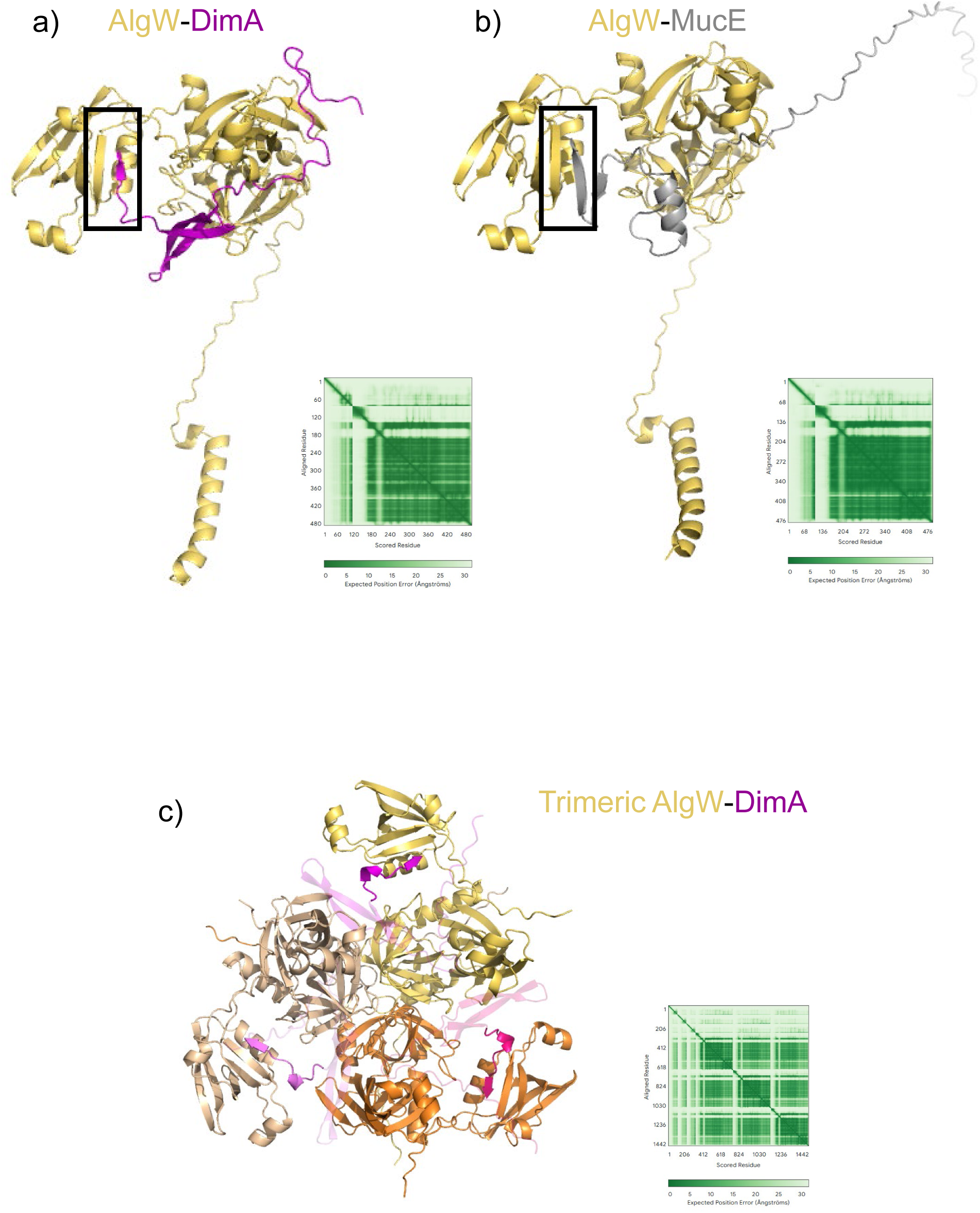
Alphafold3 prediction model of AlgW-DimA resembles AlgW-MucE interaction. a-c) Alphafold3 prediction structures. Confidence in inter- and intra-chain positioning is visualized using the Predicted Aligned Error (PAE) plot (bottom right). In the PAE heatmap, each cell represents the expected positional error (in Ångströms), dark green indicates low expected positional error (high confidence), while light green to white indicates higher uncertainty. a) Multimer model for AlgW (in yellow) and DimA (in purple), the interaction between the DimA WTF motif and AlgW PDZ domain has been boxed. b) Multimer model for AlgW (in yellow) and MucE (in grey), the interaction between the MucE WVF motif and AlgW PDZ domain has been boxed. c) Multimer model for trimeric AlgW (in yellow, wheat, and orange) and three DimA copies (in magenta, pink, and purple). To highlight the position of the C-terminal region of DimA in the structure, the last 10 amino acids are shown with 0% transparency (fully opaque), while the remainder of the structure is displayed at 70% transparency.

**Supplemental Figure 8.**
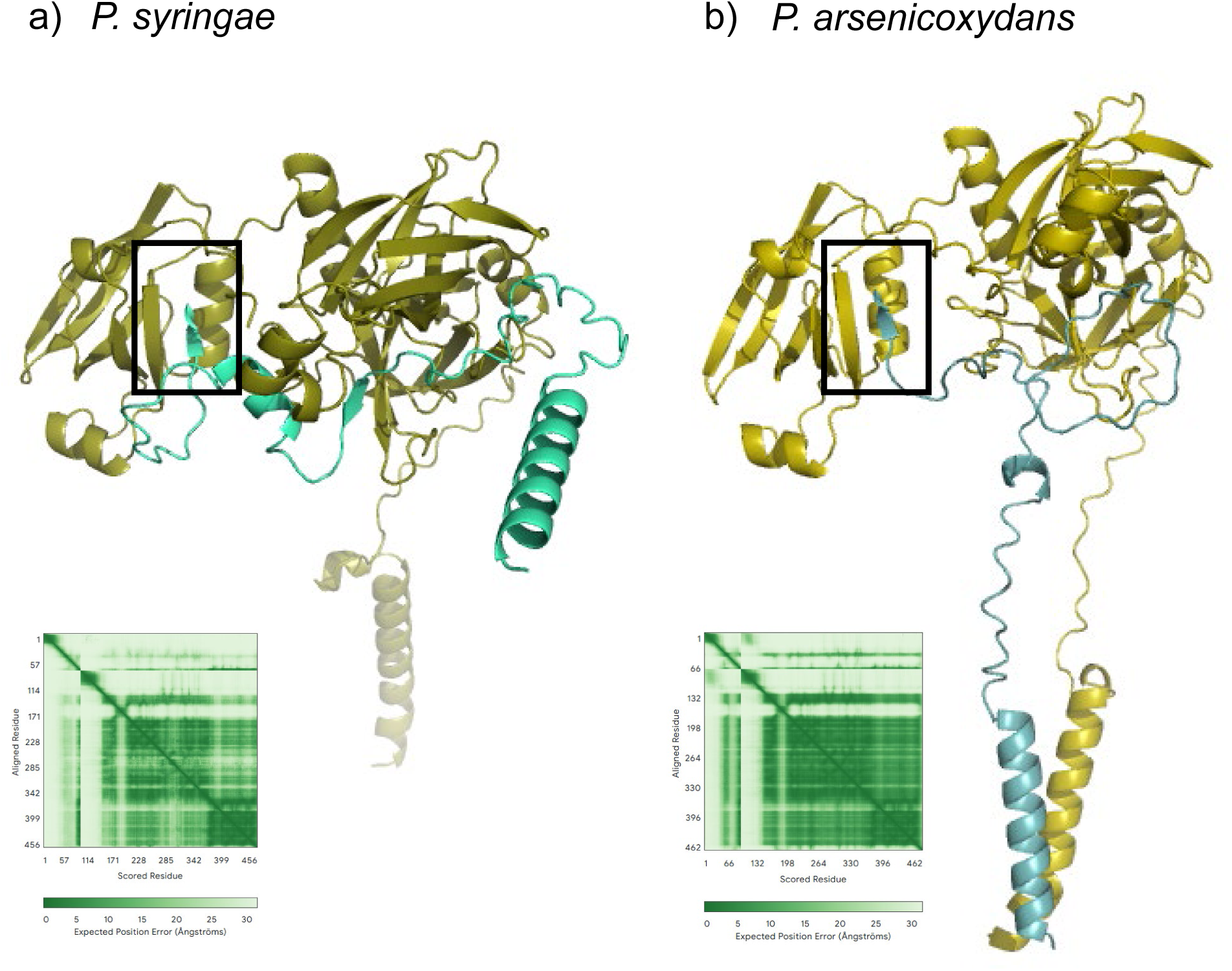
Alphafold3 prediction model of AlgW-DimA homologues in other *Pseudomonas species.* a-b) Alphafold3 prediction structures. Confidence in inter- and intra-chain positioning is visualized using the Predicted Aligned Error (PAE) plot (bottom left). In the PAE heatmap, each cell represents the expected positional error (in Ångströms), dark green indicates low expected positional error (high confidence), while light green to white indicates higher uncertainty. a) Multimer model for AlgW (olive) and DimA (turquoise) homologs in *Pseudomonas syringae*, the interaction between the DimA WVF motif and AlgW PDZ domain has been boxed. b) Multimer model for AlgW (yellow) and DimA (cyan) homologs in *Pseudomonas arsenicoxydans*, the interaction between the DimA LVF motif and AlgW PDZ domain has been boxed.

**Supplemental Figure 9.**
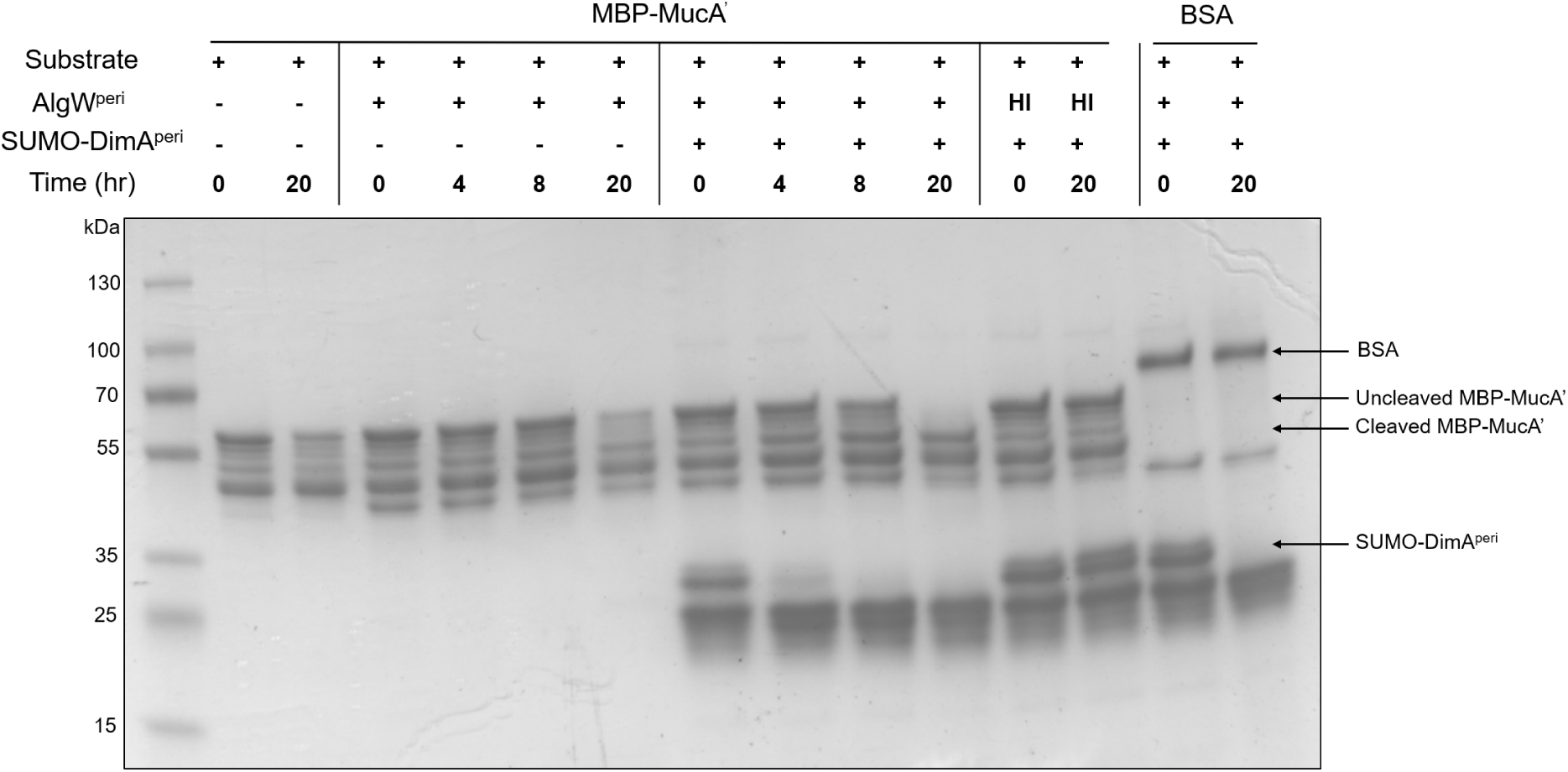
DimA is a specific activator for the AlgW-mediated proteolysis of MucA. SDS-PAGE gel stained with Coomassie Brilliant Blue for the i*n vitro* analysis of MBP-‘MucA proteolysis in the presence of AlgW protease and the DimA activator. Molecular weight markers (kDa) are shown on the left. MBP-‘MucA was purified as a mixture consisting primarily of full-length MBP-‘MucA and MBP, along with faint bands corresponding to likely truncated MBP-‘MucA intermediates.

**Supplemental Figure 10.**
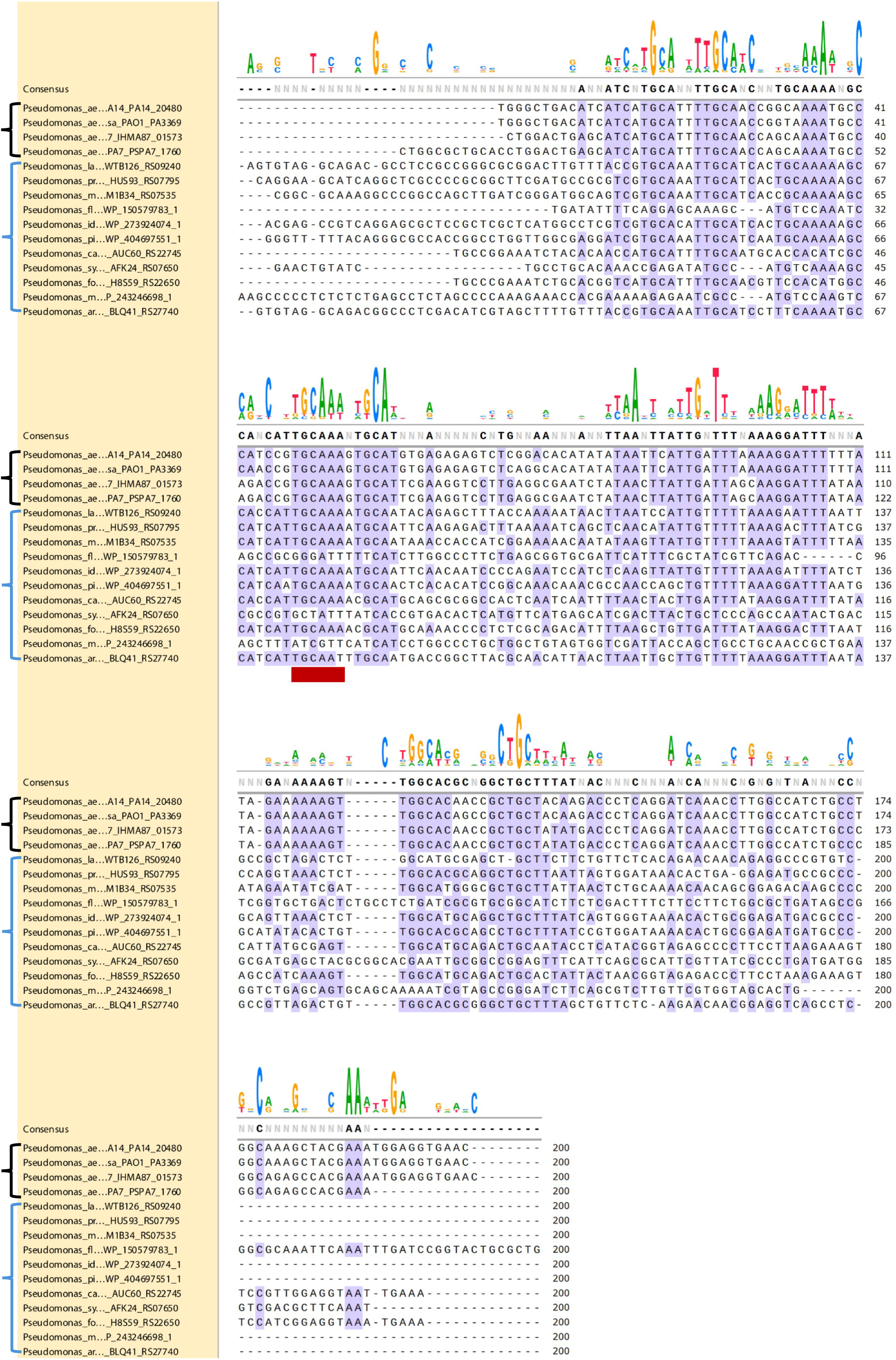
AlgB binding site in the *dimA* promoter region is conserved among *Pseudomonas* species. Nucleotide sequence alignment of the *dimA* promoter region from representative *Pseudomonas aeruginosa* isolates (black cluster) and other *Pseudomonas* species (blue cluster). The analyzed region spans from -200 to -1 bp upstream of the translation start site. The red bar indicates the proposed AlgB binding DNA motif ([37]). Conserved residues (≥50% identity threshold) are highlighted in purple. Alignment was performed using Clustal Omega and visualized with SnapGene.

**Supplemental Figure 11.**
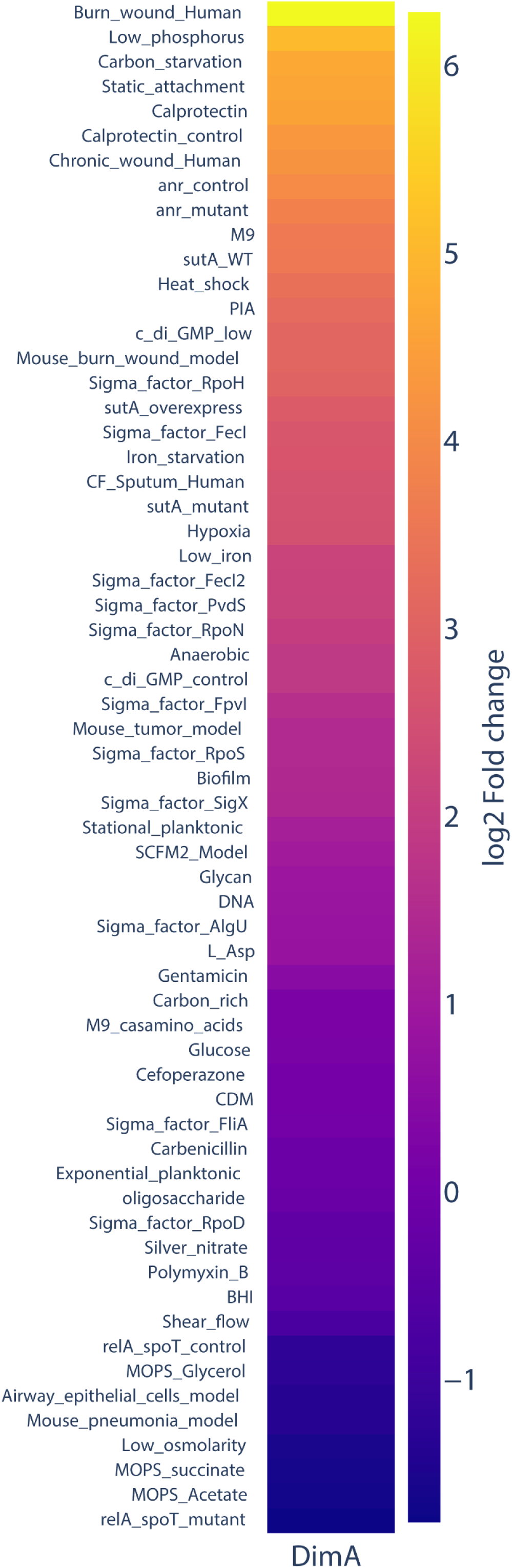
d*i*mA expression is elevated in surface-associated and stress conditions in publicly available *Pseudomonas aeruginosa* transcriptomes. Heatmap showing the log₂ fold change in expression of the dimA gene across publicly available RNA-seq datasets, relative to LB-grown P. aeruginosa UCBPP-PA14. Conditions are ranked from highest to lowest dimA induction. RNA-seq data were downloaded from the NCBI Sequence Read Archive (SRA) and aligned to the P. aeruginosa PA14 genome (assembly accession: GCA_000014625.1) using a custom nf-core-compatible pipeline.

## SUPPLEMENTAL TABLES

**TABLE S1:** Transposon insertion locations.

| PA14 ID<sup>a</sup> | Gene name and description of encoded protein |
| --- | --- |
| PA14_03340 | hypothetical protein |
| PA14_10700 | *bphP*, bacteriophytochrome |
| PA14_13490 | hypothetical protein |
| PA14_17140 | *mucP*, membrane-associated zinc metalloprotease |
| PA14_20480 | hypothetical protein |
| PA14_36420 | probable sensor/response regulator hybrid |
| PA14_41230 | *clpX*, ATP-dependent protease ATP-binding subunit ClpX |
| PA14_41240 | *clpP*, ATP-dependent Clp protease proteolytic subunit |
| PA14_44430 | hypothetical protein |
| PA14_48160 | probable sensor/response regulator hybrid |
| PA14_54430 | *algU*, RNA polymerase sigma factor AlgU |
| PA14_57760 | *algW*, AlgW protease |
| PA14_57940 | *rpoN*, RNA polymerase factor sigma-54 |
| PA14_63910 | *cntI*, putative nicotianamine synthase |
| PA14_66400 | conserved hypothetical protein |
| PA14_72380 | *algB*, two-component response regulator |
| PA14_72410 | hypothetical protein |

a: annotation from www.pseudomonas.com ([71])

**TABLE S2:**
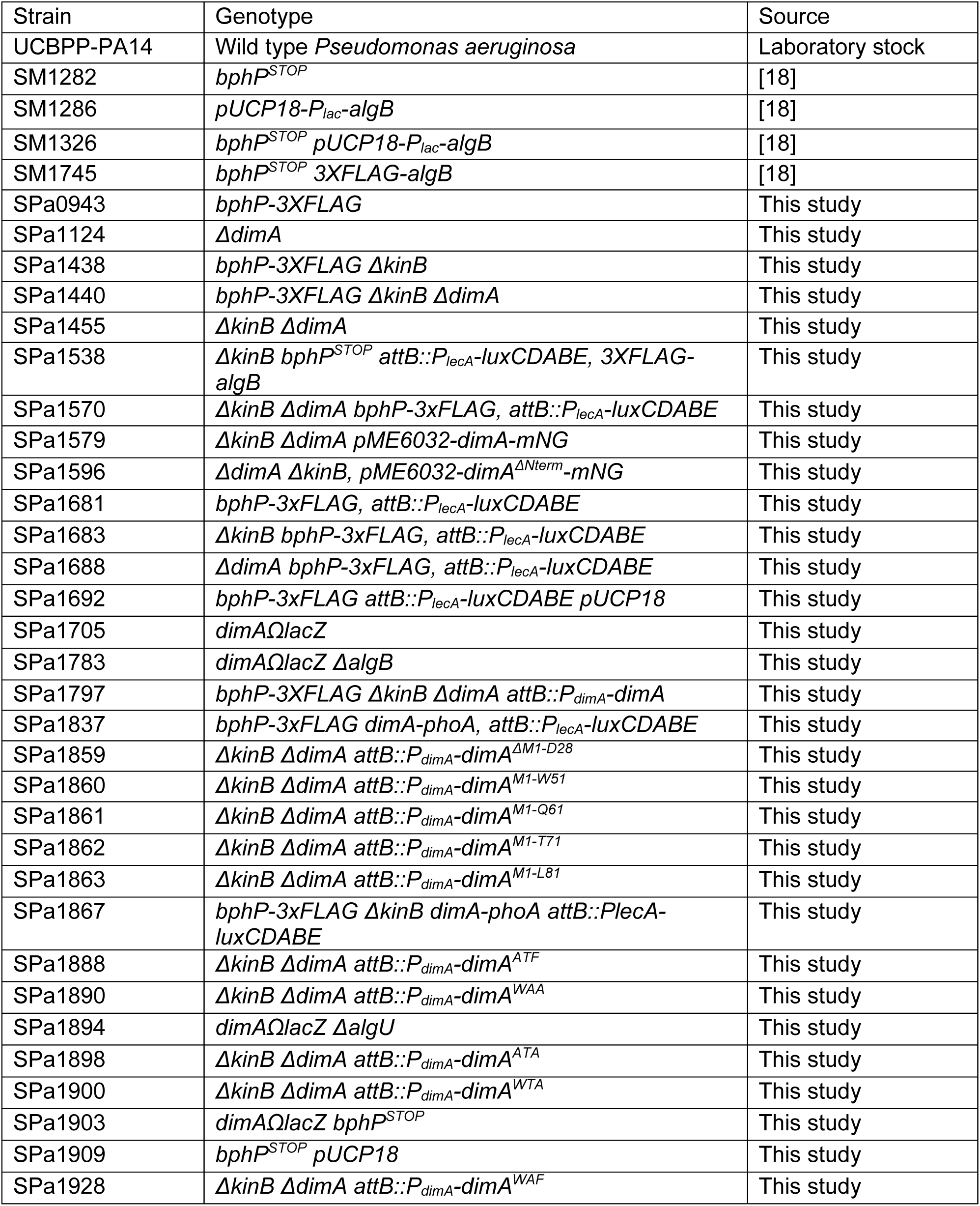
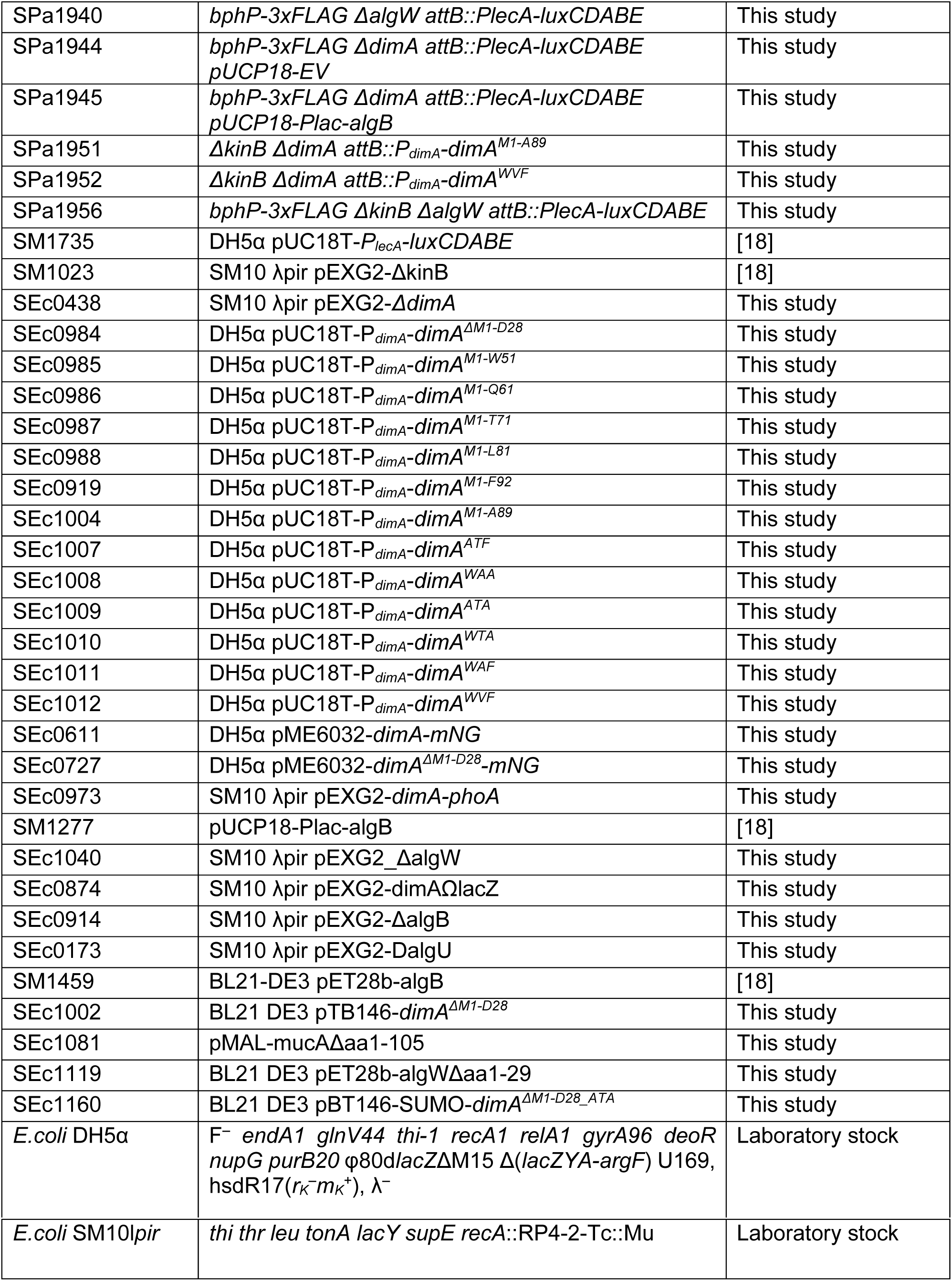
Bacterial strains.

**TABLE S3:**
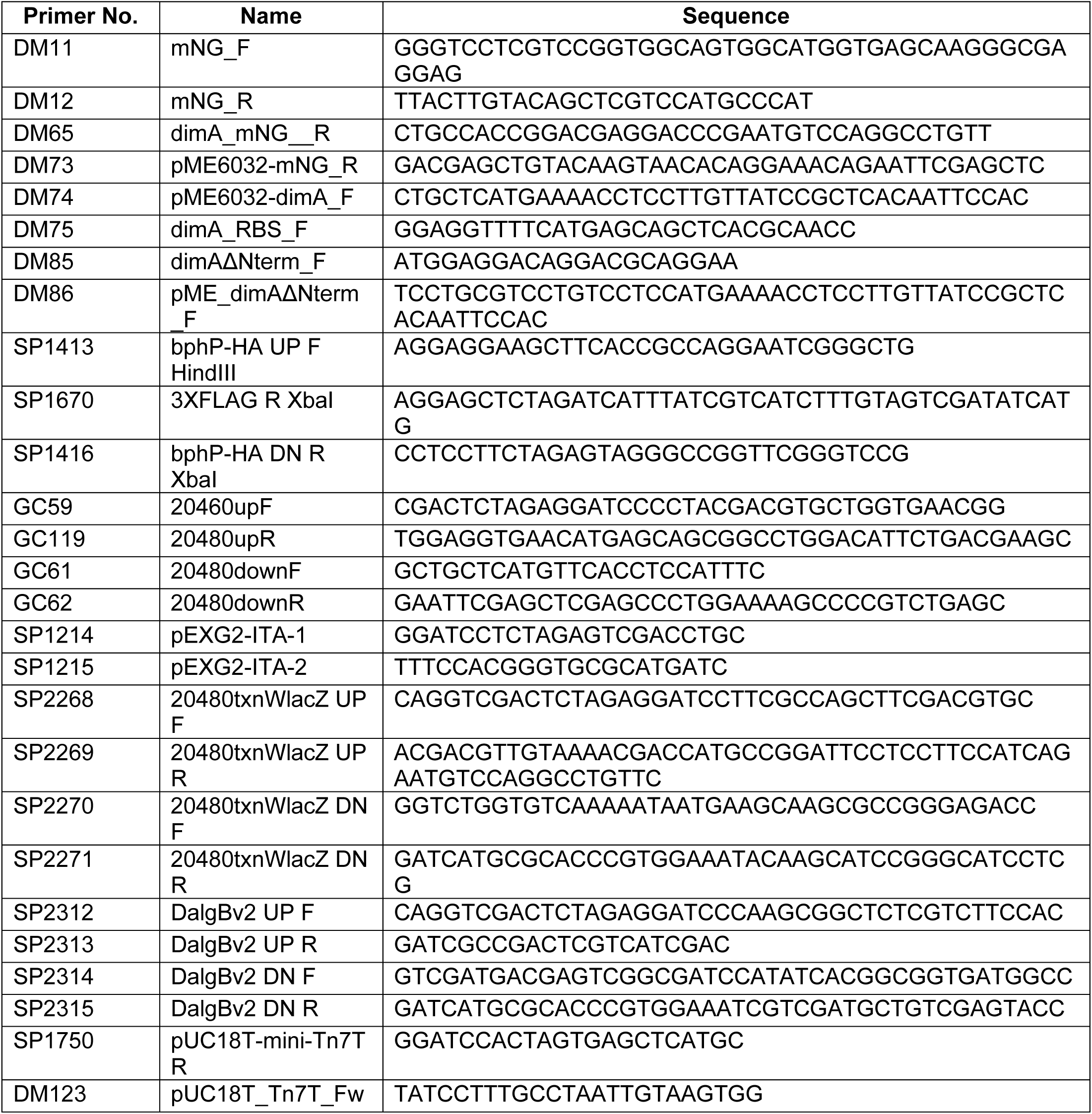
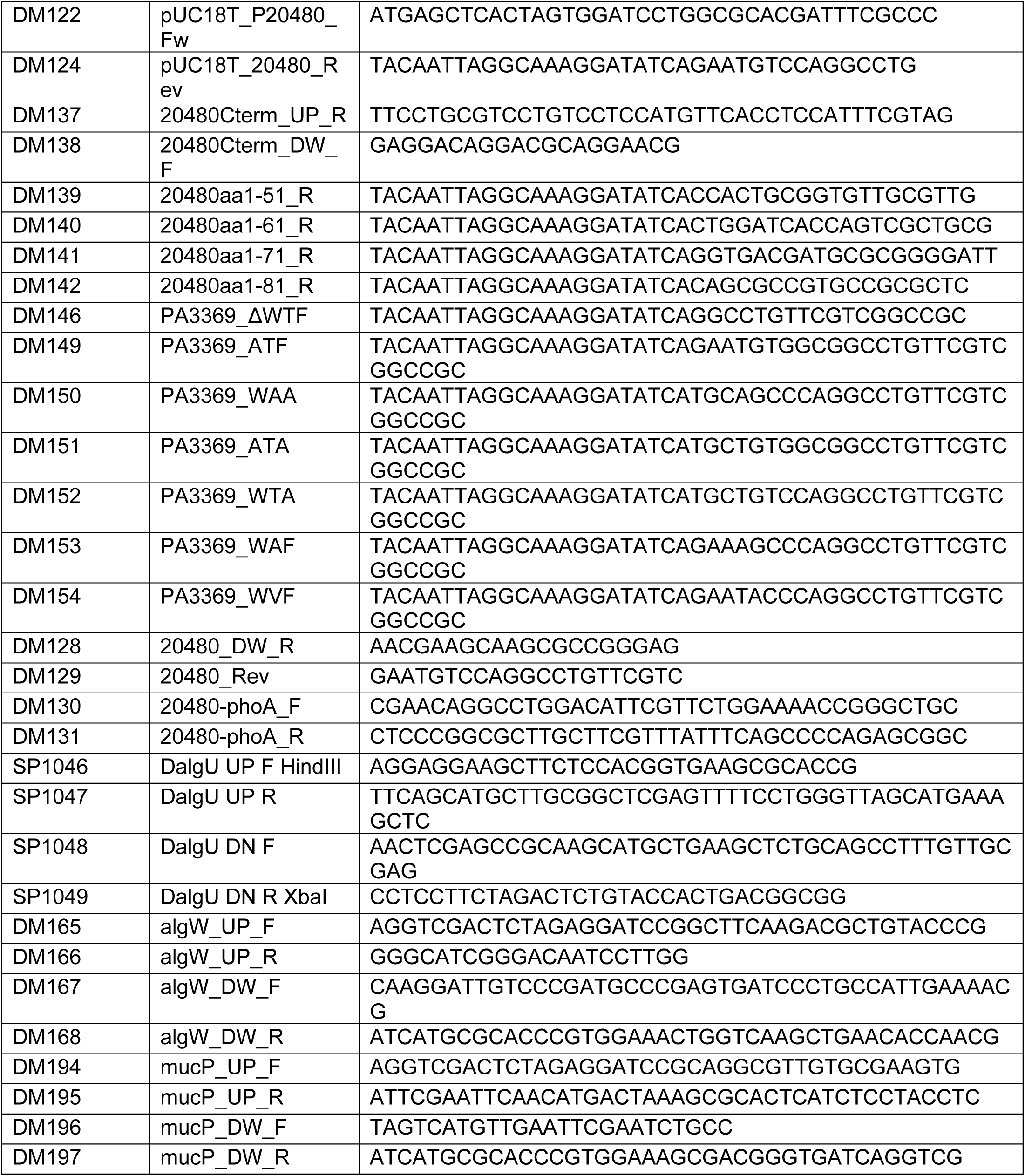
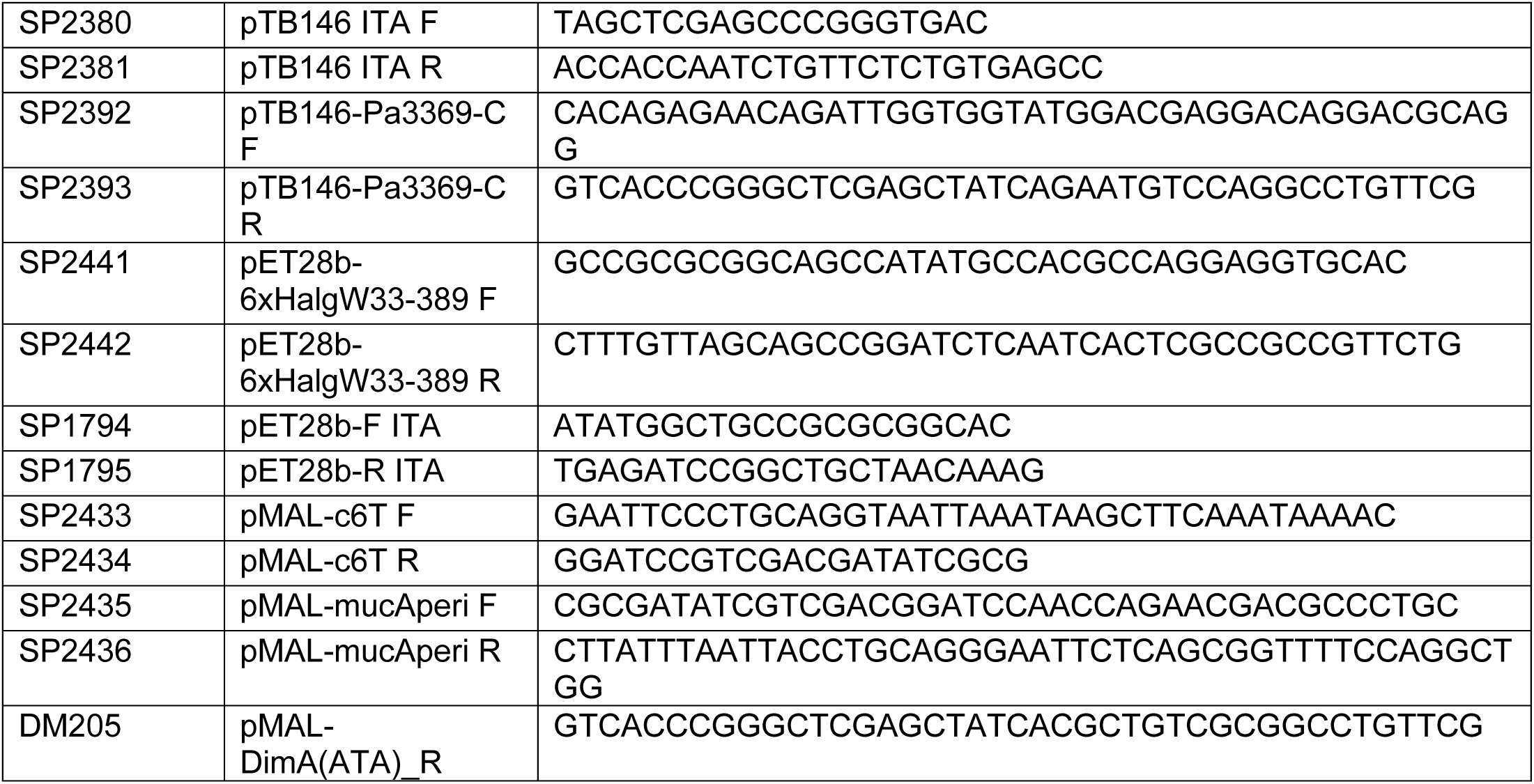
Primer List.

## Notes

### Competing Interest Statement

The authors have declared no competing interest.

